# Balance between autophagy and apoptosis determines anoikis resistance: a mathematical model on cell fate

**DOI:** 10.1101/2025.06.09.658583

**Authors:** Sayoni Maiti, Adithya Chedere, Mohit Kumar Jolly, Nagasuma Chandra, Annapoorni Rangarajan

**Affiliations:** Interdisciplinary Programme in Mathematical Sciences, IISc Mathematics Initiative, Indian Institute of Science, Bengaluru, India; Department of Biochemistry, Indian Institute of Science, Bengaluru, India; Department of Bioengineering, Indian Institute of Science, Bengaluru, India; Department of Developmental Biology and Genetics, Indian Institute of Science, Bengaluru, India

**Keywords:** Metastasis, anoikis resistance, AMPK, Akt, ERK, autophagy, apoptosis, mathematical modeling

## Abstract

Although several cancer cells are shed into circulation, most die due to stress, including loss of matrix attachment which triggers anoikis. A subset, however, acquires anoikis resistance, and survives to seed metastasis. How matrix-deprived cancer cells balance autophagy and apoptosis to decide cell fate remains unclear. Here, we formulated a deterministic ordinary differential equation (ODE)-based model of Akt-AMPK-ERK signal interaction to study this balance. Simulations revealed that high AMPK activity, with heterogenous ERK and low pAkt levels promote survival, whereas high pAkt shifts towards apoptosis. Molecular perturbation analyses revealed a signaling hierarchy in which Akt dominates over AMPK which further dominates over ERK. This model further predicts that the extent of autophagy determines apoptotic outcomes with high autophagy favoring survival of matrix-detached MDA-MB-231 cells. Together, our study provides a mathematical framework linking signaling dynamics to cell-fate decisions governing anoikis resistance.

**Graphical Abstract:** 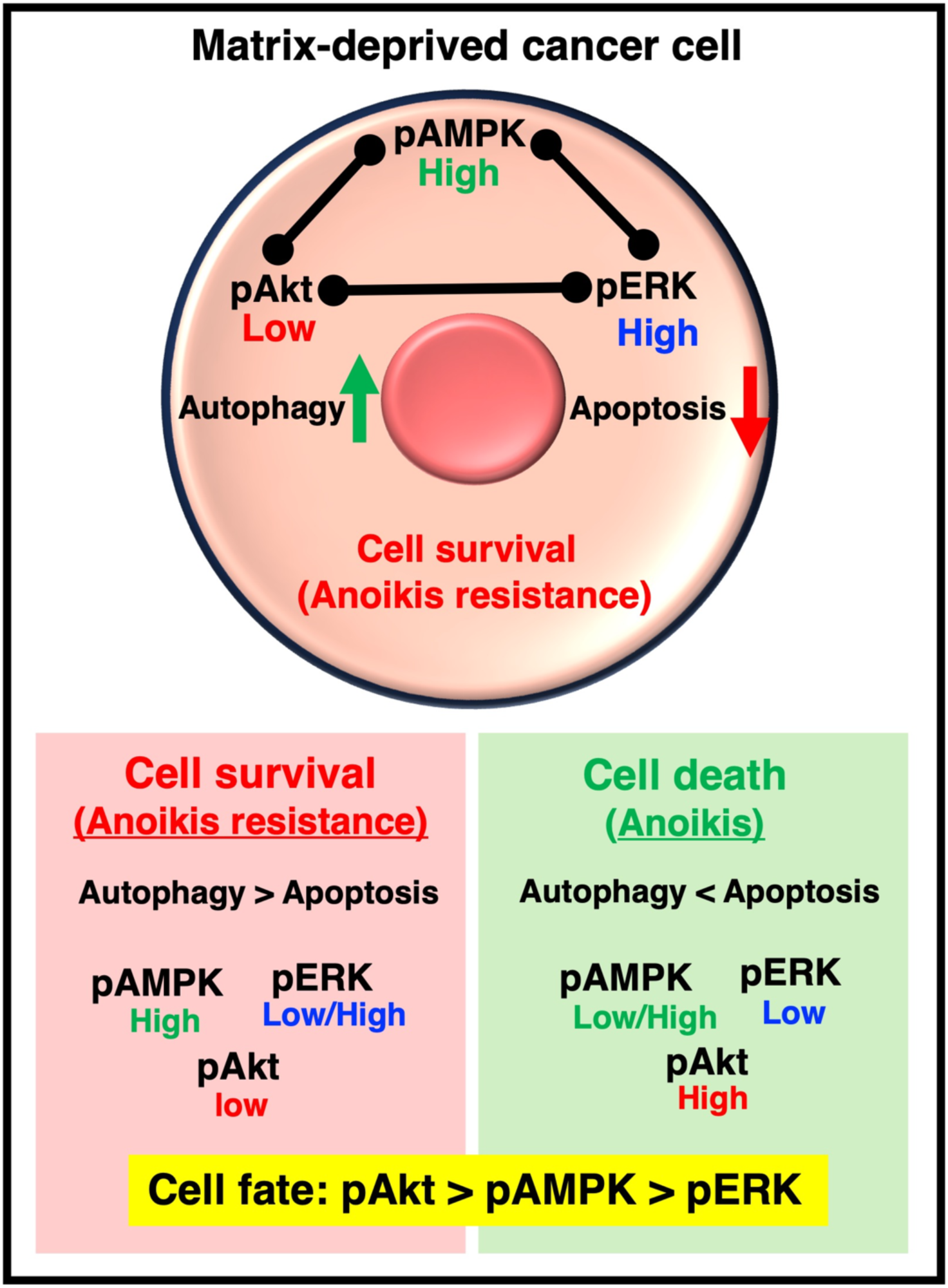

## INTRODUCTION

Metastasis is one of the primary causes of cancer-related mortality in most forms of solid tumors. It is a multistep biological process by which cancer cells detach from the primary site, intravasate into the vasculature, survive in the circulation, extravasate at a secondary site and develop into a secondary tumor.^1^ While all of the above steps are crucial to seed metastasis, the ability of the epithelial cancer cell to survive in an anchorage-independent manner is a critical step. As a barrier to developing metastases, cancer cells usually undergo apoptosis due to detachment from their extracellular matrix (ECM) by a process coined as ‘‘anoikis”.^2^ Thus, tumor cells that acquire malignant potential have developed mechanisms to resist anoikis and thereby survive after detachment from their primary site and seed metastasis by a mechanism known as anoikis resistance.^3^

Cancer cells acquire the resistance to anoikis by altering several signaling and metabolic pathways.^4,5^ A few signaling pathways known to regulate the cellular stress induced by matrix deprivation is along the Akt-AMPK-ERK axis.^6^ Protein kinase B or Akt is another serine-threonine protein kinase that has emerged as a crucial regulator of widely divergent cellular processes including apoptosis, proliferation, differentiation, and metabolism.^7^ Although Akt has been extensively studied in solid tumors, its role in matrix-detached or migrating cancer cells is rather controversial. Some reports suggest that Akt activation is one of the most common mechanisms to achieve anoikis resistance in cancer cells.^8^ However, work from our lab showed that there is decrease in Akt activity in matrix-deprived cancer cells in multiple cell lines.^6^ AMP-activated protein kinase (AMPK) is a serine-threonine protein kinase that is a central regulator of cellular energy homeostasis and is known to get activated upon multiple bioenergetic stresses, such as, hypoxia, glucose deprivation, and matrix deprivation.^9,10^ The RAS/RAF/MEK/ERK (Extracellular signal Regulated Kinase) pathway is a highly conserved signaling pathway which connects extracellular signals from cell surface receptors to machinery that regulates multiple critical physiological processes, including growth, proliferation, differentiation, migration, and apoptosis.^11^ ERK is shown to get activated in matrix-deprived state as compared to the matrix-attached state.^12^

In addition to the intricate interplay of signaling pathways, anoikis resistance is also influenced by the balance between cell survival and cell death mechanisms which decides the fate of the cell. Numerous literatures have suggested that cell fate can be studied as a balance between autophagy and apoptosis.^13,14^ Autophagy is the process by which cells degrade and recycle proteins and organelles to maintain intracellular homeostasis^15^ and is also activated upon matrix detachment.^16,17^ Generally, autophagy plays a protective role in cells, but disruption of autophagy mechanisms or excessive autophagic flux usually leads to cell death.^15^ On the other hand, apoptosis is a cellular mechanism that is responsible for programmed cell death and is also activated upon matrix detachment.^18^

Prior work from our laboratory has shown that there is increase in the levels of active AMPK, ERK and decrease in the levels of active Akt upon matrix deprivation as compared to the matrix-attached state.^6,12^ We have also shown that there exists a double-negative feedback loop between AMPK and Akt in both matrix-attached and matrix-deprived condition.^6,19^ Interestingly, though both pAMPK and pERK are found to be high in matrix-deprived condition, we have shown that upon AMPK inhibition by compound C, there is increase in ERK activity^12^, and upon ERK inhibition by PD98059 there is increase in AMPK activity^12^, suggesting a double-negative feedback loop between AMPK and ERK as well. Further, AMPK phosphorylates and activates PEA15, which prevents the activation of ERK.^20^ ERK has been shown to phosphorylate and inhibit LKB1, an upstream activator of AMPK, and thereby blocking the activation of AMPK by LKB1 in BRAF(V600E)-driven melanoma.^21^ These indicate the existence of a double-negative feedback loop between AMPK and ERK in matrix-deprived condition. Though the Akt-ERK axis has not been explored enough in the matrix-detached state, literature shows that treating matrix-attached lung cancer cells with inhibitors of ERK signaling increased Akt phosphorylation while inhibition of Akt signaling increased ERK phosphorylation.^22^ Literature also shows that MEK inhibitor caused the induction in Akt activity in HER2-positive breast cancer^23^, while ERK inhibition resulted in sustained Akt phosphorylation under the stress of low glucose in HEK 293 cells^24^, and phosphorylation of Raf by Akt led to inhibition of ERK in MCF7 cell line.^25^ The above evidence indicates that there may exist a double-negative feedback between Akt and ERK as well.

Upon matrix deprivation, there is initiation of autophagy, but interestingly, the completion of autophagy is halted, and there is increase in apoptosis.^12^ In addition to the interesting crosstalks that exist among Akt, AMPK and ERK in matrix-attached or matrix-detached condition, each of these three proteins influence cell fate pathways like autophagy and apoptosis. We have observed that hyperactivation of Akt enhanced the levels of active caspase-3 and reduced the number of anchorage independent colonies in matrix-deprived condition.^6^ This indicates that Akt impairs cell survival and enhances apoptosis of matrix-deprived cells. Secondly, we have observed that AMPK inhibition enhanced the levels of active caspase-3 and reduced the number of anchorage-independent colonies in matrix-deprived condition.^6^ This indicates that AMPK promotes cell survival and halts apoptosis of matrix-deprived cells. Thirdly, we have shown that ERK inhibition reduces the levels of active casepase-3 and increases autophagy completion in matrix-detached cells.^12^ This indicates that ERK halts autophagy completion and facilitates apoptosis in matrix-deprived condition. Although these data have independently highlighted Akt-AMPK and AMPK-Erk cross talks in regulating apoptosis and autophagy, yet little is known about how the interplay among these three kinases – Akt, AMPK and ERK – regulates the cell fate mechanisms like apoptosis and autophagy, and how does the fine balance between these mechanisms contributes towards the anoikis resistance of matrix-deprived MDA-MB-231 breast cancer cells.

This study focuses on deciphering how cancer cells decide their fate (autophagy or apoptosis) under matrix deprivation using mathematical modelling. We integrated existing knowledge on anoikis resistance into a rigorous analytical framework and formulated an ordinary differential equation (ODE)-based mathematical model on cell fate pathways (autophagy and apoptosis) influenced by Akt, AMPK and ERK. Our dynamical model is built using existing experimental data of signaling pathways (Akt, AMPK and ERK) and cell fate pathways (autophagy and apoptosis) in metastatic breast cancer cells (MDA-MB-231) subjected to matrix deprivation (summarized in **Table S1**). Molecular perturbation of key proteins depicted that though low levels of pERK support cell survival, but this may happen so until levels of pAkt are maintained low. At extreme low levels of pERK, there is a possibility of high levels of pAkt which may dominate over AMPK in matrix-deprived state and favor anoikis. Overall, this work provided multiple insights on the molecular interplay among key kinases Akt, AMPK and ERK and their effects on cell-fate decision of apoptosis and autophagy. This model also depicted that, based on the extent of autophagy (which is prominently induced in this system), apoptosis is either high or low. Hence, for anoikis-resistant MDA-MB-231 cells, the model predicts that cell fate decisions are influenced by the balance between autophagy and apoptosis, majorly influenced by survival pathways like autophagy.

## RESULTS

### Majority of MDA-MB-231 cells survive upon matrix deprivation and show enhanced autophagy

Detachment from the matrix triggers cell death by apoptosis, termed as anoikis. To counter cell death, matrix detachement also triggers autophagy, a stress response that promotes cell survival. To begin to understand how matrix-deprived cancer cells regulate the balance between autophagy and apoptosis to overcome the stress of matrix deprivation, we first investigated the status of apoptosis and autophagy in MDA-MB-231 metastatic breast cancer cells subjected to matrix deprivation for 24 h. MDA-MB-231 cells grown in matrix-attached (att) condition depicted spindle-shaped morphology **(Figure S1A)** and grown in matrix-deprived (sus) condition showed the formation of large aggregates **(Figure S1B)**. To study the apoptosis profile of MDA-MB-231 cells upon matrix deprivation, we performed AnnexinV/PI apoptosis assay. AnnexinV bind phosphatidylserine (PS) located on the outer surface of the plasma membrane and marks the early apoptotic cells. PI is impermeant to live cells and early apoptotic cells, but stains dead cells with red fluorescence, binding tightly to the nucleic acids in the cell.^26^ After staining a cell population with Annexin V and PI, the percentage of live cells were scored as Annexin V(-)/PI(-), necrotic cells were detected by Annexin V(-)/PI(+), early apoptotic cells were detected by Annexin V(+)/PI(-) and late apoptotic cells were detected by Annexin V(+)/PI(+). The percentage of late-apoptotic cells in att condition was 3.8% **(Q2, Figure 1A)** and in sus condition was 9.1% (**Q2, Figure 1B**). The percentage of live cells in att condition was 94.8% **(Q3, Figure 1A)** and sus condition was 88.9% **(Q3, Figure 1B)**. The statistical analysis (**Figure 1C**) also depicted that there is no significant difference among the cell populations of att versus sus condition. These data indicate that a majority of MDA-MB-231 cells are anoikis-resistant and survive upon matrix deprivation albeit a small population succumbs to anoikis. This trend has also been shown in the literature.^27,28^

**Figure 1:**
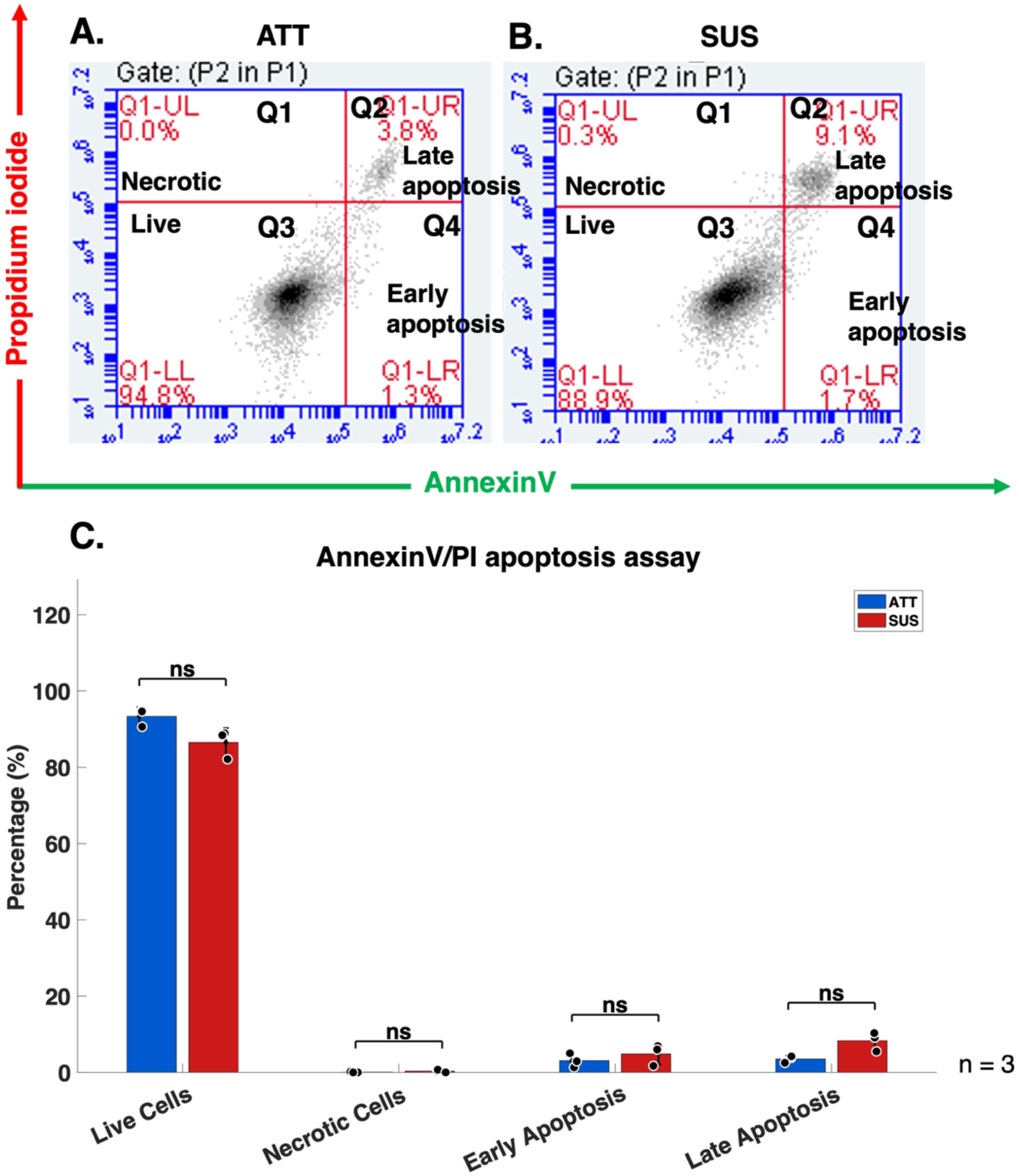
Majority of MDA-MB-231 cells survive upon matrix deprivation. Representative flow cytometry dot plots of apoptosis of MDA-MB-231 cells exposed to **(A)** matrix-attached state (Att) and **(B)** matrix-deprived state (Sus) for 24 h. The lower left quadrant (Q3) shows percentage of live cells [Annexin V(-)/PI(-)], upper left quadrant (Q1) shows percentage of necrotic cells [Annexin V(-)/PI(+)], lower right quadrant (Q4) shows percentage of early apoptotic cells [Annexin V(+)/PI(-)] and upper right quadrant (Q2) shows percentage of late apoptotic cells [Annexin V(+)/PI(+)], n=3. **(C)** Graph represents the quantification of all four cell populations. Error bars represent SEM, n=3, *P < 0.05; **P < 0.01; *** P < 0.001; ns: not significant.

To study the balance between autophagy and apoptosis in MDA-MB-231 cells upon matrix deprivation, we performed a dual-staining experiment where cells were stained with both active caspase 3/7 reagent (written as caspase henceforth) and autophagosome detection reagent (written as autophagosome henceforth). For controls, we stained the cells with caspase and autophagosome reagents, individually.

**Figure 2A-F** shows the flow cytometry dot plots of matrix-attached (Att; upper panels) and matrix-deprived (Sus, lower panel) MDA-MB-231 cells. Each dot plot consists of four quadrants: unstained, caspase only, autophagosome only, and both. To study the active-caspase expression profile, MDA-MB-231 were stained with active caspase 3/7 reagent and analysed by Fluorescence-assisted cell sorting (FACS). **Figure 2A** shows that 65% (yellow) of the adherent cells are do not show caspase expression (unstained), and 7% (red) of the cells depict high caspase expression in the Att cells. **Figure 2B** shows that upon suspension, 56% (yellow) of the cells do not show caspase expression (unstained), and 10% (red) of the cells depict high caspase expression. The above data depicts that high active 3/7 caspase expression is shown by 7% (Att) - 10% (Sus) of the population that corroborates the above data on AnnexinV/PI apoptosis assay (Att: 4%, Sus: 9%). The remaining percentage of cells (white) express low levels of active caspase 3/7. Multiple evidences show the presence of low levels of active caspase 3/7 in healthy cells and report survival from caspase activation after treatment with drugs or radiation.^29^

**Figure 2:**
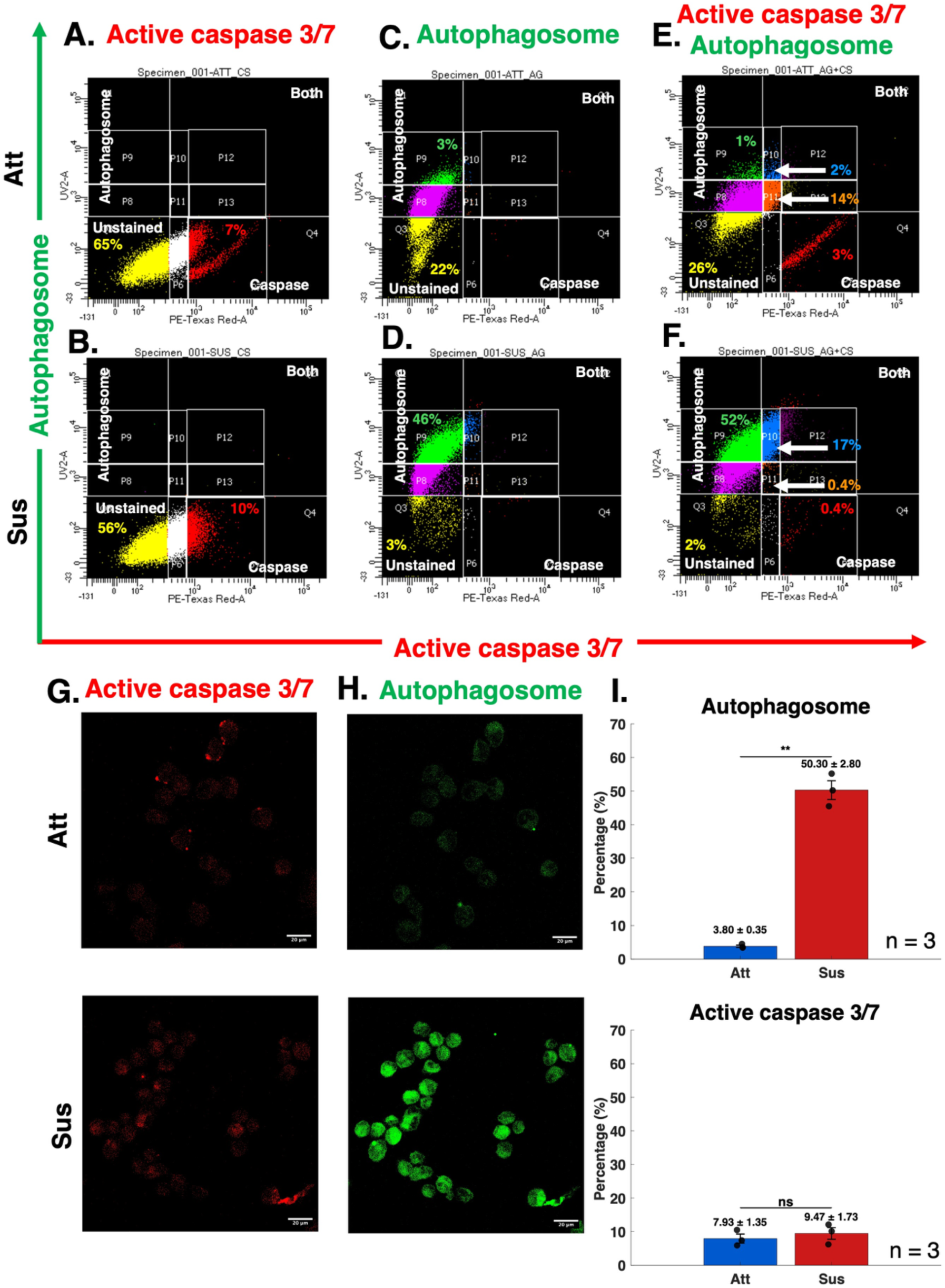
Majority of MDA-MB-231 cells show enhanced autophagy upon matrix deprivation. Active caspase 3/7 and autophagosome expression in matrix-deprived condition: Fluorescence-activated cell sorting was done for MDA-MB-231 cultured in attached and suspension condition for 24 h, stained with **(A-B)** active caspase 3/7 reagent and **(C-D)** autophagosome detection reagent, individually and **(E-F)** simultaneously. X-axis depicts the expression of active caspase 3/7 and Y-axis depicts the expression of autophagosome, n=3. Photomicrographs show representative fluorescence images of **(G)** active caspase 3/7 (red), **(H)** autophagosome (green) in MDA-MB-231 cells cultured in attached (Att) or suspension (Sus) condition for 24 h. Representative fluorescent images were visualized with confocal microscopy (Z-stack, scale bar, 20 μM, magnification: 40x). **(I)** Graphs represent the quantification of percentage of cells expressing autophagosome (top panel) and active caspase 3/7 (bottom panel). Error bars represent SEM, n=3,*P < 0.05; **P < 0.01; *** P < 0.001; ns: not significant.

To study the autophagosome expression profile, MDA-MB-231 were stained with autophagosome detection reagent and analysed by Fluorescence-assisted cell sorting (FACS). **Figure 2C** shows that 22% (yellow) of the adherent cells do not show autophagosome expression (unstained) while 3% (green) of the cells depict high autophagosome expression. The remaining percentage (pink) accounts for the basal autophagy in the cells. **Figure 2D** shows that upon suspension, 3% (yellow) of the cells do not show autophagosome expression (unstained) while 46% (green) of the cells depict high autophagosome expression. The above data depict that matrix-deprived MDA-MB-231 cells exhibit enhanced autophagy.

To further study the expression profile of active-caspase and autophagosome in the same cell population, MDA-MB-231 were stained with active caspase 3/7 and autophagosome detection simultaneously and analysed by Fluorescence-assisted cell sorting (FACS). The expression low caspase-low autophagosome profile (orange) is 14% for Att cells and 0.4% for Sus cells (**Figure 2E**). The expression of low caspase-high autophagosome profile (light-blue) is 2% for Att cells and 17% for Sus cells (**Figure 2F**). This data indicates that majority of the cells with low caspase expression (**Figure 2A-B**, white) have shifted to the autophagosome positive regions (**Figure 2E-F**, orange and light blue). Literature has also shown that Caspase 3 and caspase 7 promote cytoprotective autophagy during non-lethal stress conditions in human breast cancer cells.^30^

Post FACS sorting, the caspase-positive, autophagosome-positive, and dual-positive (caspase-autophagosome-positive) cells were collected and imaged by confocal microscopy. We observed the presence of active-caspase in both Att and Sus groups and the expression appeared to be similar across both the groups (**Figure 2G**). The expression of autophagosome was much higher in Sus cells as compared to the att condition (**Figure 2H**). The statistical analyses also depicted a clear increase in the expression of autophagosome (**Figure 2I; top panel**) whereas, there is no significant difference in the levels of active-caspase (**Figure 2I; top panel**) in Sus cells as compared to att cells.

Overall, the above data indicate that there is enhanced autophagy in matrix-deprived MDA-MB-231 cells, and this could be the reason for suppressed cell death in these cells. And indeed these values of autophagy and survival have been emphasized in the model and its predictions.

This intrigued us to question how is the balance between autophagy and apoptosis decided in matrix-deprived cancer cells to overcome the stress due to matrix deprivation and enable survival. To answer this question, we utilized mathematical modeling techniques to formulate a mathematical model which could capture the above biology in additional to crucial signaling events that are altered in these conditions.

### Construction and calibration of anoikis-resistance mathematical model

Multiple pathways including AMPK/ACC, calcium/ROS signaling, integrin signaling, MAPK/ERK, HIF1, PEA15, NFκB/Wnt, Akt/mTOR/S6K1, anabolism/catabolsim and autophagy/apoptosis regulate anoikis and are perturbed upon matrix deprivation^4,6,31,32^ as summarized in **Figure 3Ai**. Though the above signaling events are highly interlinked, there exits a sequence in which the cue of matrix deprivation is transferred from one level to another. Upon matrix deprivation, the growth factor and integrin signaling are perturbed which effect RAS/RAF/MEK pathways. Additionally, there is a spike in cytosolic calcium and ROS signaling that activates AMPK via LKB1 and CaMKKβ. Akt-AMPK-ERK are unknown to influence eachother and regulate anoikis. Further downstream are mTOR/S6K1, PEA15, NFκB and HIF1 which regulate metabolism and autophagy/apoptosis. Previously, we built a mathematical model that captured the attachment-detachment signaling and linked the effects of AMPK-Akt-mTOR-S6K1 to metabolism.^33^ In this work, we focus on Akt-AMPK-ERK and their effect on autophagy and apoptosis. To reduce the complexity of the above system, we chose a few proteins or secondary messengers to formulate a mathematical model with minimum number of variables needed to study anoikis resistance. To create a mathematical model that captures the sequence of events during matrix deprivation (starting from the cue of detachment to the cell fate of survial/death), we coarse-grained the extended system (**Figure 3Ai**) into a minimum network consisting of Akt, AMPK and ERK signaling nodes coupled to cell fate pathways like autophagy and apoptosis (**Figure 3Aii**) based on experimental data summarized in **Table S1**. This is named the ‘anoikis resistance model’, which is a deterministic model wherein, protein-protein interactions are described by ordinary differential equations and outputs report temporally variable protein concentration. The model consists of eight variables: Matrix, cytosolic calcium (Ccal), phospho-Akt (pAkt), phospho-AMPK pAMPK, phospho-ERK (pERK), phagophore, autophagosome, active-caspase (aCaspase) which capture the dynamics of attachment-detachment signaling. Phagophore and autophagosome are variables associated with autophagy and aCaspase is associated with apoptosis. All the variables are listed in **Table S2,** initial conditions on all the variables are listed in **Table S3** and all the parameters are listed in **Table S4**. All the equations and their brief descriptions are also presented in the **Supplemental Information** section.

**Figure 3:**
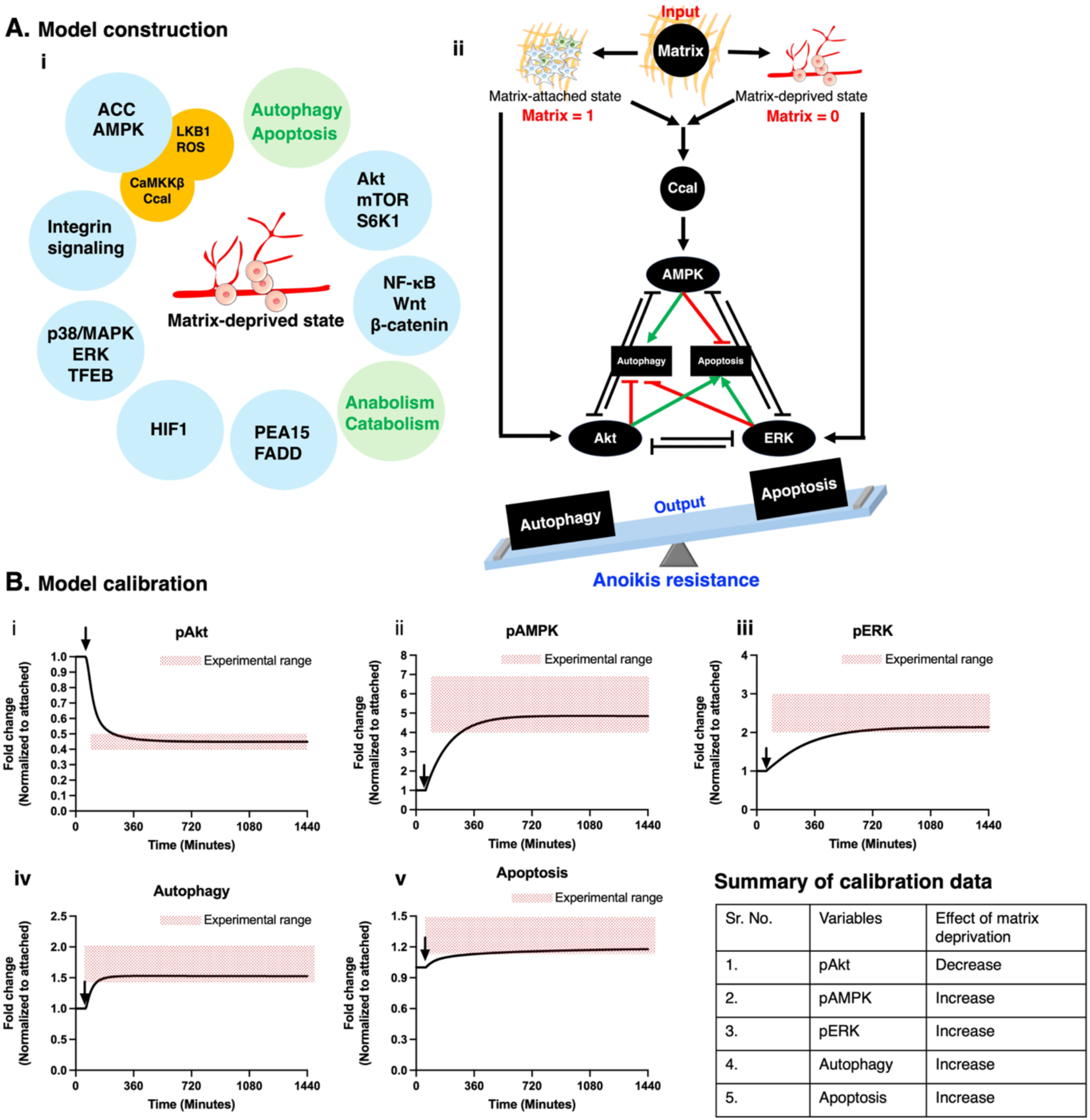
Anoikis resistance mathematical model. (A) Model construction. **i** Multiple pathways that regulate anoikis and are perturbed upon matrix deprivation. **ii** The anoikis resistance mathematical model depicting the model input: matrix, model players: Akt, AMPK and ERK, and model outputs: apoptosis and autophagy. **(B) Model calibration.** The calibration set consists of temporal dynamics of simulated concentration of following protein species in MDA-MB-231 cell line for 24 h period following matrix deprivation: **i** pAkt, **ii** pAMPK, **iii** pERK, **iv** Autophagy, and **v** Apoptosis. The above calibration data is summarized in the table. The red shared area depicts the range of experimental data. The black solid line depicts the temporal dynamics predicted by the model. The black arrow indicates the time point of induction of matrix deprivation. X axis depicts time in minutes, where, first 60 minutes is the matrix-attached state and remaining is the matrix-deprived state. Y axis depicts the fold change normalized to matrix-attached condition.

The following assumptions have been made while building the model:

1. A double-negative feedback loop between Akt and ERK in matrix-deprived state.
2. The total levels of Akt, AMPK, ERK and proCaspase were considered to be 10 nM.
3. Since autophagosome are formed from phagophores and phagophore are dynamic structures, the amount of phagophore is written in the form of an ODE instead of a constant value.
4. Apoptosis is studied by the levels of aCaspase. Once effector caspases are activated, the cell is programed to die. The accumulation of cleaved executioner caspases 3/7 is often considered a hallmark of an irreversible commitment to apoptosis and the cleavage-specific antibodies that detect the active forms of caspase 3/7 are used to identify apoptotic cells.^34^ Biochemically, the name ‘caspase’ is a contraction of cysteine-dependent aspartate-specific protease. Their enzymatic properties are governed by a dominant specificity for protein substrates containing aspartate and by the use of a cysteine side chain for catalyzing peptide bond cleavage. This proteolytic event is irreversible.^35^ Hence, the levels of active or cleaved caspases do not revert back to pro-caspase forms. Hence, in the equation for aCaspase, only the activation of caspase has been included. There are no terms included for the reduction in the levels of aCaspase. Therefore, the levels of aCaspase keep increasing with time, while all other variables attain steady state.
5. Since multiple players are involved in Akt-AMPK-ERK signaling pathways and how they influence autophagy and apoptosis, all their interactions are captured as hill functions.

The first assumption on the double-negative feedback loop between Akt and ERK in the matrix-deprived deprived state has been experimentally tested as shown in **Figure S2.** Results show increase in the levels of pAkt upon ERK inhibition in matrix-deprived condition. Investigating the reverse (effect of Akt inhibition on the levels of pERK) is challenging to perform in matrix-deprived condition owing to the extremely low levels of pAkt in the matrix-deprived state. Hence, the reverse feedback was assumed based on the experimental data available for matrix-attached condition.^22^

The input to the anoikis resistance model is the presence or absence of extracellular matrix or simply ‘Matrix’. For the matrix-attached state, the value of Matrix is assigned as 1, which denotes the presense of matrix attachment. For the matrix-detached state, the value of Matrix is assigned as 0, which denotes the absence of matrix attachment. The key model players are Akt, AMPK and ERK. In the matrix-attached state, growth factor and integrin (attachment-mediated) signaling activates Akt. Active Akt inhibits AMPK and ERK. Whereas, in the matrix-deprived state, AMPK is activated via CAMKKβ due to the spike in cytosolic calcium (Ccal). Matrix detachment also leads to ERK activation. Detachment-triggered AMPK inhibits Akt and ERK. Active ERK also inhibits AMPK and Akt. Hence, multiple double-negative feedback loop exists among AMPK, Akt and ERK as depicted in **Figure 3Aii**.

In addition to the interaction of AMPK, Akt and ERK among themselves, these proteins also influence autophagy and apoptosis. AMPK activation promotes autophagy whereas AMPK inhibition enhanced caspase-3 activity and reduced anchorage independent colonies.^6^ Hence, AMPK promotes survival mechanisms like autophagy and inhibits apoptosis. Though Akt levels are low in matrix-deprived state, hyperactivation of Akt enhanced caspase-3 activity and reduced anchorage independent colonies.^6^ Hence, Akt inhibits autophagy but promotes apoptosis in matrix-deprived cancer cells.^6^ ERK inhibition reduced apoptosis and enhanced autophagy completion.^12^ Therefore, ERK promotes apoptosis and inhibits autophagy. The experimental data depicting the above points are shown in the **Table S1**.

Though excess autophagy can lead to cell death, in this model we use the formation of autophagosome as a surrogate for autophagy induction (as used previously shown^13^) and portray autophagy as a mechaism of cell survival. The formation of autophagosome indicates that the cell is in survival mode. The activation of caspase-3 is used as a surrogate for apoptosis or cell death. In this model, we define survival as autophagy outweighing apoptosis. This scenario also exemplifies anoikis resistance. Alternately, we define cell death as apoptosis outweighing autophagy. This scenario exemplifies anoikis. The above description is pictorially depicted in **Figure 3Aii**.

Model calibration was done to fit the mathematical model to the experimental data. This calibration was done by retrieving some model parameters from the literature and estimating the remaining parameters to get the best fit of the model to the experimental data. All the experimental data used for model calibration are depicted in **Figure S3** and **Figure S4** and tabulated in **Table S1A.** The results of model calibration are depicted in **Figure 3B**. The results show the dynamics of pAkt (**Figure 3Bi**), pAMPK (**Figure 3Bii**) and pERK (**Figure 3Biii**) consistent with experimental observations under matrix deprivation. The model also captured the increase in autophagy (**Figure 3Biv**), and apoptosis (**Figure 3Bv**) upon matrix deprivation. Additionally, model parameters were tweaked to capture a couple of experimental data from activation/inhibition studies. Model captured the decrease in pAkt upon enhancing the levels of pAMPK in the matrix-attached state (**Figure S5A**), the increase in the levels of pAkt upon ERK inhibition (**Figure S5B**) and increase in the levels of pERK upon Akt inhibition (**Figure S5C)**.

### Mathematical analyses of anoikis resistance mathematical model

Following the model calibration, we performed multiple analyses on the model to evaluate the steady state of the system and to study the stability of these steady states.

To evaluate the steady states of the system, the model was simulated for 24 h and beyond. The temporal dynamics of the variables for 1440 min (24 h) along with the steady state values attained in matrix-attached and matrix-deprived conditions are depicted in **Figure 4A**. Since the key model players comprise of pAkt, pAMPK and pERK, further analyses were done on the three model players. When model was simulated for longer duration, the same steady state values were obtained for pAkt, pAMPK and pERK (data not shown).

**Figure 4:**
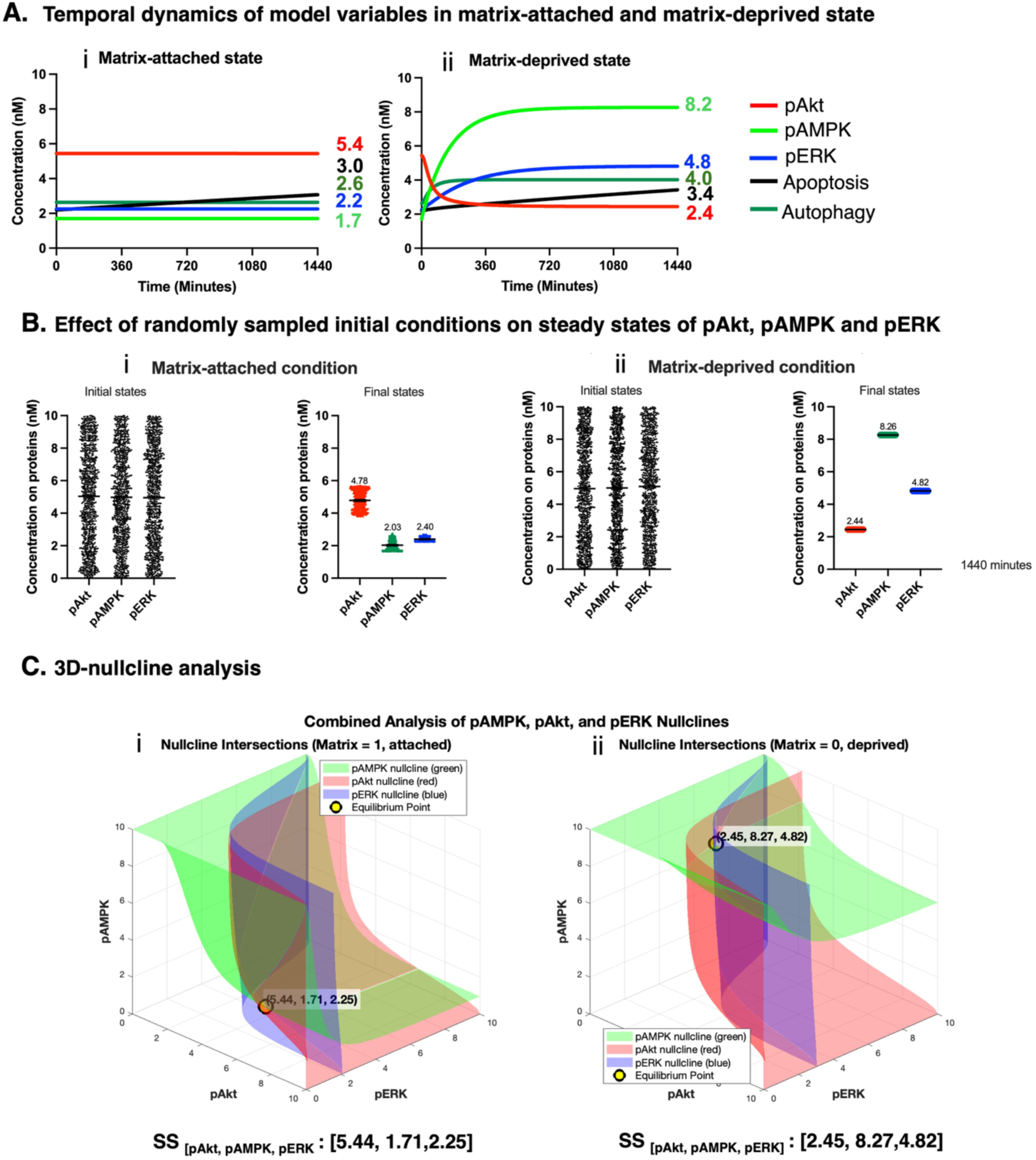
Mathematical analysis of anoikis resistance model. **(A)** Temporal dynamics of model variables in **(i)** matrix attached and **(ii)** matrix-deprived state. X axis depicts time in minutes. Y axis depicts the concentration of pAkt, pAMPK and pERK in nM or levels of apoptosis and autophagy. **(B)** Effect of randomly-sampled initial conditions on steady states of pAkt, AMPK and pERK in **(i)** matrix attached and **(ii)** matrix-deprived state. For each matrix-attached and matrix-deprived state, the panel of the left shows 1000 random initial states which were sampled from a uniform distribution of range 1-10 nM and the panel on the right shows the final steady state values at 1440 min, with the mean value depicted on the graph. X axis depicts the three model players (pAkt, pAMPK and pERK). Y axis depicts the concentration of these proteins in nM. **(C)** 3D-nullcline analysis for **(i)** matrix attached and **(ii)** matrix-deprived state. The red surface shows pAkt nullcline (d/dt(pAkt) = 0; X axis). The green surface shows pAMPK nullcline (d/dt(pAMPK) = 0;) The blue surface shows pERK nullcline (d/dt(pERK) = 0). The intersection point of three nullclines depict the stable steady state value of the **(i)** matrix-attached state [5.44, 1.71, 2.25] and **(ii)** matrix-deprived state [2.45, 8.27, 4.82].

To study the effect of initial conditions on the final steady state values of these model players, the initial conditions were randomly-sampled, and the model was simulated in both the matrix-attached and matrix-deprived conditions. Results show that irrespective of the value of initial condition of pAkt, pAMPK or pERK, the steady state values of these proteins were 4.78, 2.03 and 2.40 respectively in the matrix-attached state (**Figure 4Bi**) and the steady state values of these proteins were 2.44, 8.26 and 4.82 respectively, in the matrix-deprived state (**Figure 4Bii**), when simulated for a total time duration of 1440 min (24 h). The above analysis was also repeated for longer duration and similar results were obtained (data not shown). These results depicted that the model is robust and captures the biology irrespective of any initial conditions.

To confirm the steady states of the system, 3D-nullclines for pAkt, pAMPK and pERK were obtained by plotting 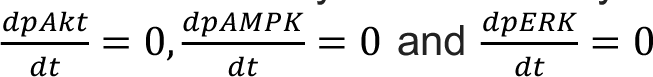 and [inine], (**Figure S7**) in a three-dimensional space with axes pAkt (X-axis), pAMPK (Y-axis) and pERK (Z-axis). **Figure 4Ci** shows the intersection of pAkt, pAMPK and pERK 3D-nullclines plotted simultaneously for the matrix-attached state. The single intersection point reveals the unique steady state value [pAkt: 5.44, pAMPK: 1.71, pERK: 2.25] of the system in matrix-attached state. **Figure 4Cii** shows the intersection of pAkt, pAMPK and pERK 3D-nullclines plotted simultaneously for the matrix-deprived state. The single intersection point reveals the unique steady state value [pAkt: 2.45, pAMPK: 8.27, pERK: 4.82] of the system in matrix-deprived state. Hence, this analysis confirms that the system has a single monostable steady state for matrix-attached and matrix-deprived state.

To study the stability of the steady states, steady state perturbation analysis was performed. This analysis helps to understand whether any perturbation caused at the steady state of the system affects the final steady state of the system. When the initial condition of pAkt was either decreased or increased by 50% for matrix-attached state (**Figure S8A**) or matrix-deprived state (**Figure S8D**), the temporal dynamics depicted that the system restored to original steady state values within 540 minutes. The same trend was obtained when pAMPK (**Figure S8B; Figure S8E**) and pERK (**Figure S8C; Figure S8F**) were perturbed in the same way. These results indicated that the model steady states are stable and were neither affected by changes in initial conditions nor by any perturbation to the steady state values.

Another gold standard technique to study the stability of steady states is by plotting phase portraits. **Figure S9A** depicts the phase portraits of the pAMPK 3D-nullcline in the matrix-attached state. We observe that all the vector fields point towards the steady state values of the system of matrix-attached state [pAkt: 5.44, pAMPK: 1.71, pERK: 2.25]. Likewise, the phase portraits of pAkt and pERK for matrix-attached state are shown in **Figure S9B** and **Figure S9C.** The phase portraits of pAMPK, pAkt and pERK for matrix-deprived state are shown in **Figure S9D, Figure S9E** and **Figure S9F**, respectively. We observe that all the vector fields point towards the steady state values of the system of matrix-detached state value [pAkt: 2.45, pAMPK: 8.27, pERK: 4.82]. Hence, the above results depict that the system indeed has a unique steady state for matrix-attached and matrix-deprived condition.

### Model validation

Following rigorous mathematical analyses of the calibrated model, model validation was performed by comparing simulation results against experimental data not used for model fitting previously. In the previous sections we observed that the model captured the relative change in the levels of pAkt, pAMPK and pERK while shiting from the matrix-attached to the matrix-deprived condition. This relative change is mainly because of plausible strength of the feedbacks in the network. To find out the relative strength of the feedbacks in the matrix-attached and matrix-deprived condition, we plotted a flux map. The numbers in blue pertain to the matrix-attached condition while the numbers in red pertain to the matrix-detached state. As the feedbacks and crosstalks are written in terms of Hill functions, the numbers vary between the range of 0 and 1. These hill functions are shown in **Figure 5A**. A value closer to 0 represents mild/low strength of activation or inhibition, while a value closer to 1 represents a strong/high strength of activation or inhibition. A value closer to 0.5 depicts neutral effect.

**Figure 5:**
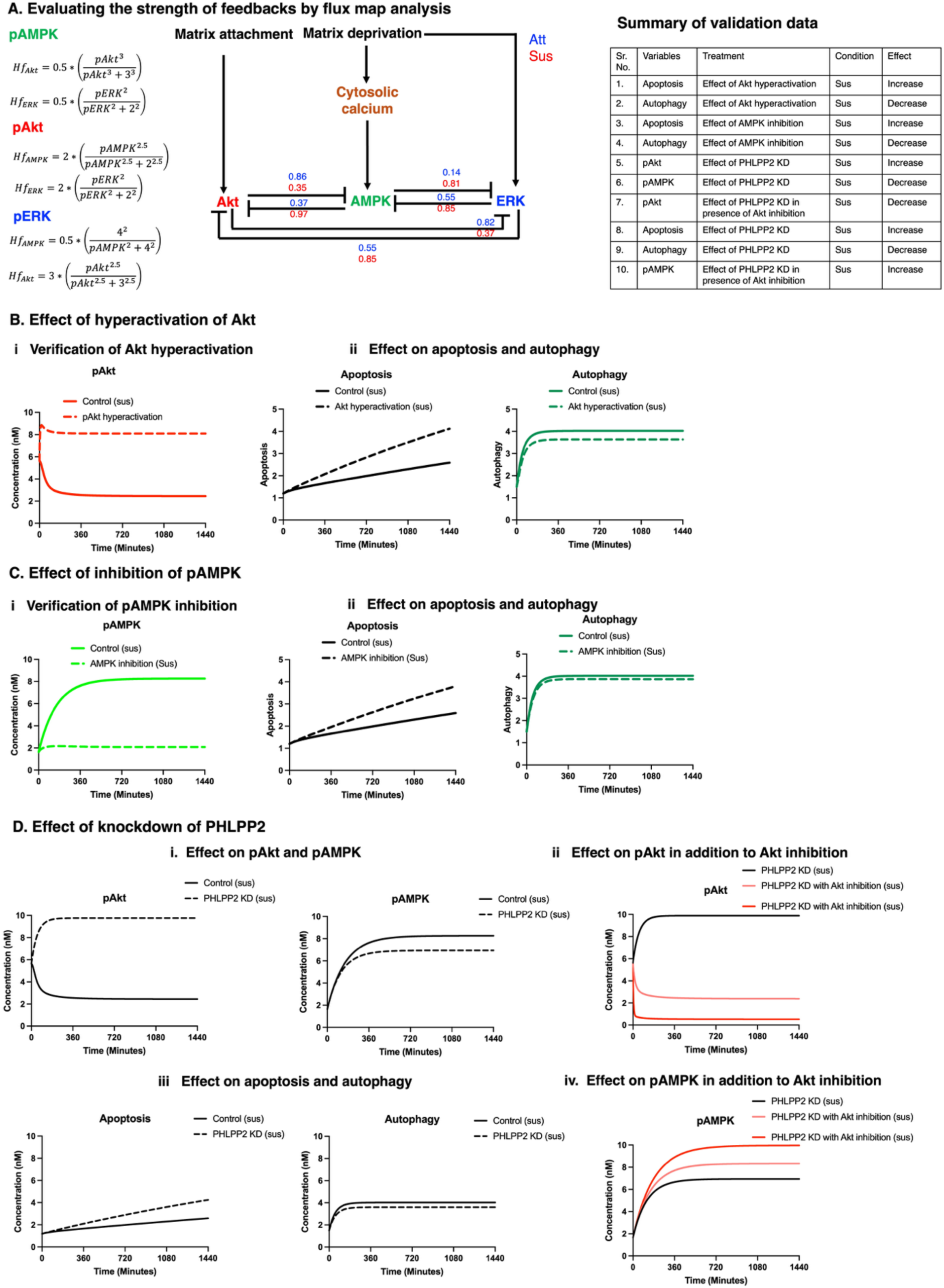
Model validation. **(A)** Flux map in matrix-attached (blue) and matrix-deprived (red) condition. Numbers along the feedbacks indicate their magnitude in terms of the corresponding hill function values (mentioned on the left), which lie between 0 and 1. The matrix-attached state is marked by high levels of pAkt and low levels of pAMPK and pERK. The matrix-deprived state is marked by low levels of pAkt and high levels pAMPK and pERK. **(B) (i)** The effect of hyperactivation of Akt on **(ii)** apoptosis and autophagy. **(C) (i)** The effect of inhibition of AMPK on **(ii)** apoptosis and autophagy. **(D)** The effect of knockdown of PHLPP2 on pAkt and pAMPK in the **(i)** absence and **(ii, iv)** presence of Akt inhibition and **(iii)** apoptosis and autophagy. X axis depicts time in minutes. Y axis depicts concentration (nM) or levels of apoptosis and autophagy.

In the matrix-attached state, integrin and growth factor signaling cause the activation of Akt. Flux map analysis in the matrix-attached state depicted that active Akt inhibited AMPK and ERK with a high strength of 0.86 and 0.82 (**Figure 5A; blue values)** respectively. However, in the matrix-deprived state, detachment from matrix activates ERK and spike in cytosolic calcium activates AMPK via CAMKKβ. Flux map analysis in the matrix-detached state depicted that active AMPK inhibited Akt and ERK with a high strength of 0.97 and 0.81 (**Figure 5A; red values**) respectively. Active ERK also inhibited AMPK and Akt with high strength of 0.85 and 0.85 (**Figure 5A; red values**). Since active Akt levels are low in matrix-deprived state, the strength of inhibition of AMPK and ERK by Akt was also low 0.35 and 0.37 (**Figure 5A; red values**), respectively. These results indicated that the strength of the feedbacks were biological relevant.

Literature shows that hyperactivation of Akt in matrix-deprived state is detrimental to the cells. It promoted anoikis by enhancing apoptosis by increased caspase-3 activity and reduced the number of anchorage independent colonies.^6^ Using the model, Akt hyperactivation was simulated in the matrix-deprived state (**Figure 5Bi**) and an increase in apoptosis was observed (**Figure 5Bii)**. Though anchorage independent colonies do not directly correspond to the levels of autophagy in the cell, but both anchorage independent colonies and autophagy indicate the extent of cell survival. The model does not show any significant difference in the levels of autophagy upon hyperactivation of Akt in matrix-deprived state (**Figure 5Bii)**. Similary, literature also shows that AMPK inhibition induces apoptosis via caspase-3 activation and reduces the number of anchorage independent colonies.^6^ Our model also predicted the increase in apoptosis (**Figure 5Cii)** upon AMPK inhibition (**Figure 5Ci)** as compared to the untreated control in the matrix-deprived state. The model did not show any significant difference in the levels of autophagy (**Figure 5Cii)** upon AMPK inhibition in matrix-deprived state.

Another validation was done using experimental data on the effect of PHLPP2 knockdown (KD) on AMPK and Akt in matrix-deprived condition. PHLPP2 is a phosphatase that mediates the inhibitory effect of AMPK on Akt.^6^ PHLPP2 KD removes this inhibitory effect. Experimentally, it was shown that PHLPP2 KD leads to decreased AMPK activity in matrix-deprived condition via Akt activation (**Figure S6A**). Hence, it was additionally shown that upon Akt inhibition in the PHLPP2 KD cells, there was a decrease in pAkt levels and an increase in pAMPK levels (**Figure S6B**). PHLPP2 KD can be simulated in silico by delinking the inhibitory effect of AMPK on Akt. In matrix-deprived condition, the strength of the inhibitory effect of AMPK on Akt was calculated to be 0.97 (details elaborated in flux map analysis), hence delinking was performed by assigning a value of 0.01 to the respective hill function. Upon simulating the PHLPP2 KD in silico (by delinking), the model predicted the increase in pAkt (**Figure 5Di**) and decrease in pAMPK (**Figure 5Di**). Upon inducing different degrees of Akt inhibition, in addition to in silico PHLPP2 KD in the matrix-deprived condition, the model predicted the decrease in pAkt (**Figure 5Dii**) and increase in pAMPK (**Figure 5Div**). Experimentally we have also observed that PHLPP2 KD impairs autophagy^6^ and enhances apoptosis.^6^ Upon simulating the effects of PHLPP2 KD on autophagy and apoptosis, the model predicted the increase in apoptosis upon PHLPP2 KD (**Figure 5Diii**); however, not much difference is noticed on autophagy ((**Figure 5Diii**).

### Identification of key parameters that impact the system behaviour

Parameter sensitivity analysis is done to understand the sensitivity of the model variables to local variations in individual parameters around the baseline values. Most parameters have negligible effect on key model variables, with sensitivity values close to zero (**Figure 6**). Each parameter is decreased by 10% and hence any change in the model variables above 10% or less than -10% is considered highly sensitive and any change from 8% to 10% or -8% to -10% is considered to be sensitive and any value in between - 8% to 8% is not sensitive. Interestingly, we notice a context-dependent (matrix-attached or matrix-deprived) difference in the sensitivity analysis.

**Figure 6:**
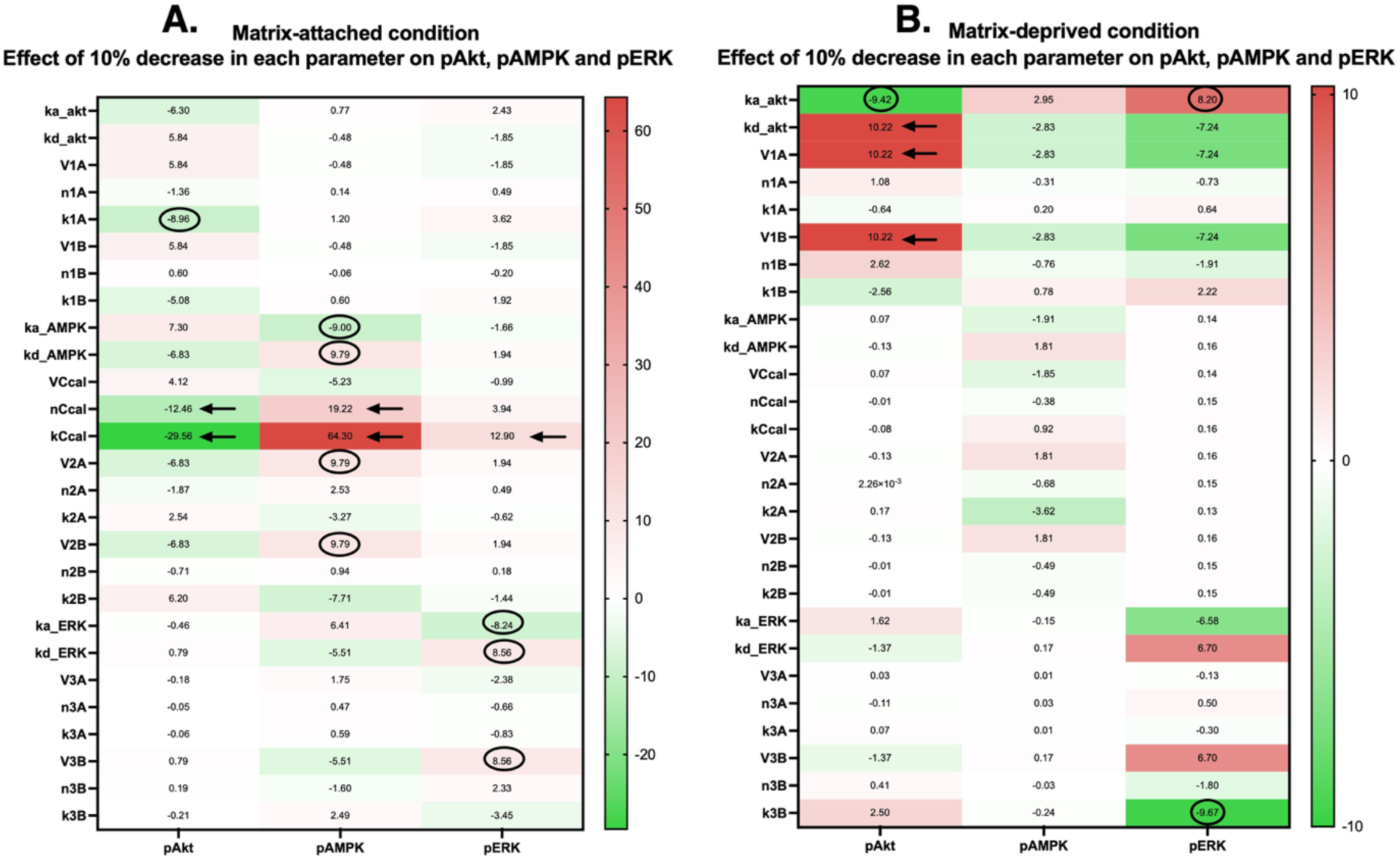
Parameter sensitivity analysis. The effect of small perturbation of all model parameters on pAkt, pAMPK and pERK in **(A)** matrix-attached state and **(B)** matrix-deprived state. Effect above 10% is shown in deep red color and less than -10% is shown in deep green color and is considered highly sensitive. Effect from from 8% to 10% is shown in light red color or -8% to -10% is shown in light green color and is considered to be sensitive and any value in between -8% to 8% is shown in white and is not sensitive.

In the matrix-attached state (**Figure 6A**), the variables (pAkt, pAMPK and pERK) are highly sensitive to kCcal (Threshold of CAMKKβ activation by Ccal). pAkt, pAMPK are sensitive to nCcal (Hill coefficient associated to Ccal). Apart from these, pAkt is sensitive to k1A (threshold of Akt inhibition by pAMPK). pAMPK is sensitive to ka_AMPK (rate of activation of AMPK), kd_AMPK (rate of deactivation of pAMPK), V2A (fold change of AMPK inhibition by pAkt) and V2B (fold change of AMPK inhibition by pERK). pERK is sensitive to ka_ERK (rate of activation of ERK), kd_ERK (rate of deactivation of pERK), and V3B (fold change of ERK inhibition by pAkt). The above results indicated that the model variables are sensitive to parameters linked to Ccal. This variable is directly linked to the input (Matrix) of the system. pAMPK and pERK are influenced by their respective rates of activation and deactivation but pAkt is not. The variables are also sensitive to parameters associated to the crosstalks between proteins.

In the matrix-deprived state (**Figure 6B**), pAkt is highly sensitive to kd_Akt (rate of deactivation of pAkt), V1A (fold change of Akt inhibition by pAMPK) and V1B (fold change of Akt inhibition by pERK) and sensitive to ka_Akt (rate of activation of Akt). pERK is sensitive to k3B (threshold of ERK inhibition by pAkt) and ka_Akt (rate of activation of Akt). These results indicated that only pAkt and pERK are sensistive to certain parameters. pAMPK appears to be rather resilient to slight changes in parameters in the matrix-deprived state.

### How do Akt-AMPK-ERK and their interplay influence the cell fate of matrix-deprived cancer cells?

Using the developed anoikis resistance model, we attempted to understand how do Akt-AMPK-ERK and their interplay impact cell fate decision pathways like apoptosis and autophagy in matrix-deprived condition. In **Figure 7**, the levels of aCaspase (black line) are used as a surrogate for apoptosis and levels of autophagosome (dark green line) are used as a surrogate for autophagy. When apoptosis outweighs autophagy, we define this as the state of anoikis in matrix-deprived cancer cells or in simple terms: cell death. When autophagy outweighs apoptosis, we define this as the state of anoikis resistance or in simple terms: cell survival. The crossover of the black and dark green lines depicts the shift from anoikis to anoikis resistance or vice versa.

**Figure 7:**
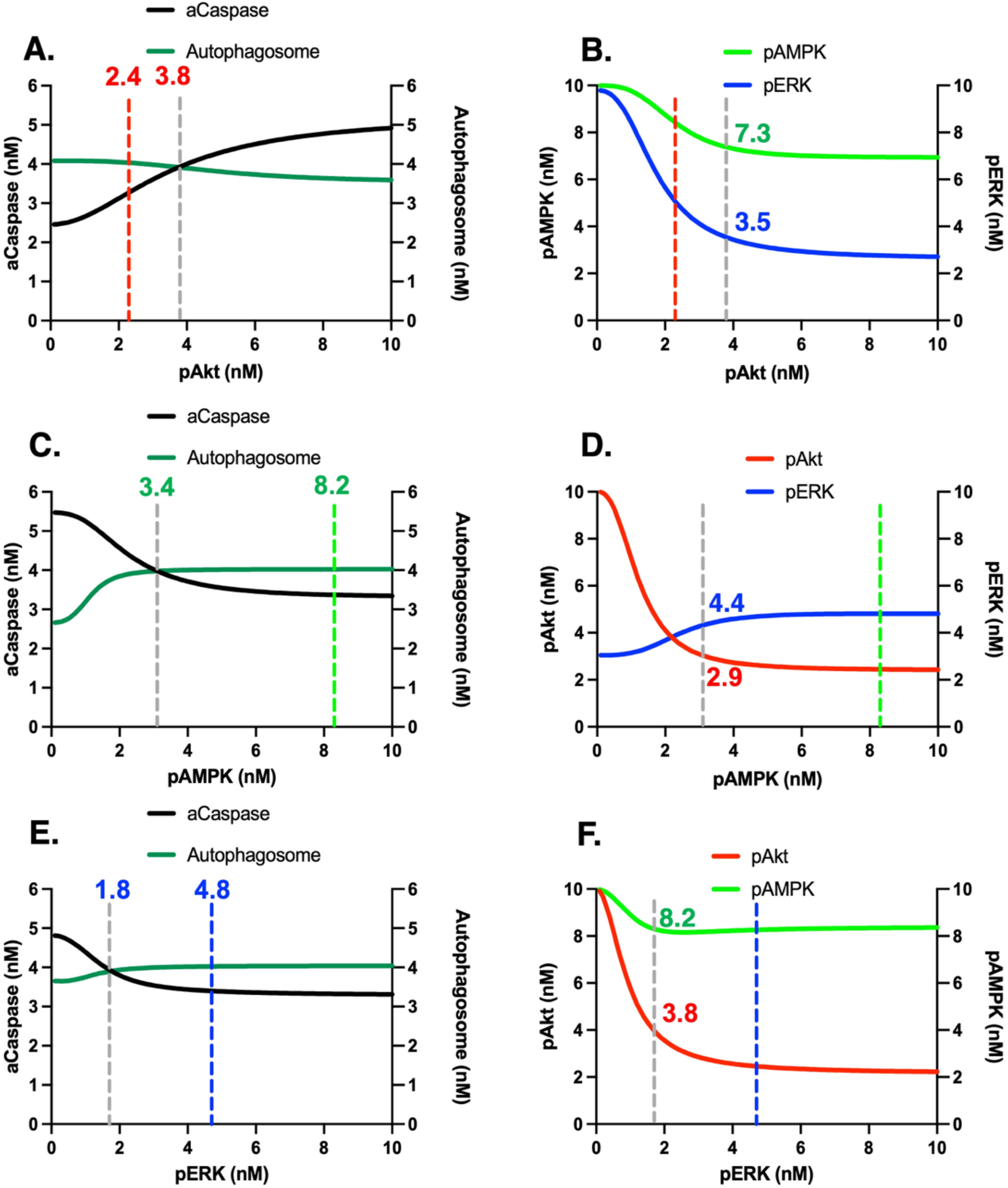
Effect of Akt-AMPK-ERK interplay on cell fate of matrix-deprived cancer cells. **(A)** The effect of differential activity Akt on apoptosis (black line) and autophagy (dark-green line). The red-dotted line depicts the concentration of pAkt at steady state and grey-dotted line depicts the concentration of pAkt at the cross over of apoptosis and autophagy. **(B)** The effect of differential activity Akt on pAMPK (light-green line) and pERK (blue line). **(C)** The effect of differential activity AMPK on apoptosis (black line) and autophagy (dark-green line). The green-dotted line depicts the concentration of pAMPK at steady state and grey-dotted line depicts the concentration of pAMPK at the cross over of apoptosis and autophagy. **(D)** The effect of differential activity AMPK on pAkt (red line) and pERK (blue line). (**E)** The effect of differential activity ERK on apoptosis (black line) and autophagy (dark-green line). The blue-dotted line depicts the concentration of pERK at steady state and grey-dotted line depicts the concentration of pERK at the cross over of apoptosis and autophagy. **(F)** The effect of differential activity ERK on pAkt (red line) and pAMPK (light-green line). X axis depicts concentration of proteins (pAkt, pAMPK, pERK). For the left panel, the left Y axis depicts the levels of apoptosis, and the right Y axis depicts

The previous analyses revealed that the steady state values of pAkt, pAMPK, and pERK in the matrix-deprived state are 2.44 nM, 8.26 nM and 4.81 nM, respectively. In this experiment, one protein is perturbed at a time (the differential equation of that protein is set to zero, and the initial condition of that protein is varied from 0 to 10 nM). Any value below its steady state value is interpreted as different levels of knockdown of that protein. Any value above the steady state value is interpreted as different degrees of overexpression of that protein. In **Figure 7**, we note the following key points:

1. The balance between apoptosis and autophagy across different concentration of the perturbed protein.
2. The concentration of the perturbed protein at the point of the crossover
3. The effect of the perturbed protein on the two other proteins
4. The levels of the other proteins at the point of the crossover. The levels are divided into three regions: 0-3 nM refers to low levels, 3-7 nM refers to intermediate levels, 7-10 nM refers to high levels.

This experiment is done specifically for the matrix-deprived condition, and each point on the graph in **Figure 7** is the steady state value at 1440 minutes (or 24 h). **Figure 7A** depicts the levels of aCaspase (left Y-axis) and autophagosomes (right Y-axis) when pAkt is varied from 0 to 10 nM (X-axis). The steady state value of pAkt (2.4 nM) in the matrix-deprived state is marked by the red-dashed vertical line. The model predicted that from 0 - 3.8 nM of pAkt, autophagy outweighs apoptosis and hence the system is resistant to anoikis. However, the shift from anoikis resistance to anoikis happens at 3.8 nM (grey-dashed vertical line) of pAkt. This also corroborates the experimental data that hyperactivation of Akt induces anoikis in matrix-deprived cancer cells.^6^ Beyond this point, apoptosis outweighs autophagy and hence the system succumbs to anoikis. This balance between apoptosis and autophagy is tightly regulated by the effect of different levels of pAkt on pAMPK and pERK. **Figure 7B** depicts the levels of pAMPK (left Y-axis) and pERK (right Y-axis) when pAkt is varied from 0 to 10 nM. The levels of pAMPK and pERK at the crossover (where pAkt is 3.8 nM) is 7.3 nM and 3.5 nM, respectively. Hence, perturbation of pAkt reveals that the region of anoikis resistance is marked by intermediate to high levels of pAkt, high levels of pAMPK and intermediate levels of pERK.

Following the perturbation of pAkt, the system was reset and pAMPK was perturbed in the same way as above. **Figure 7C** depicts the levels of aCaspase (left Y-axis) and autophagosomes (right Y-axis) when pAMPK is varied from 0 to 10 nM (X-axis). The steady state value of pAMPK (8.2 nM) in the matrix-deprived state is marked by the green-dashed vertical line. The model predicted that from 10 – 3.4 nM of pAMPK, autophagy outweighs apoptosis and the system is resistant to anoikis. However, when the levels of pAMPK reduces below 3.4 nM, apoptosis outweighs autophagy and the system succumbs to anoikis. This also corroborates the experimental data that inhibition of pAMPK favors anoikis in matrix-deprived cancer cells.^6^ **Figure 7D** depicts the levels of pAkt (left Y-axis) and pERK (right Y-axis) when pAMPK^6^ is varied from 0 to 10 nM. The levels of pAkt and pERK at the crossover (where pAMPK is 3.4 nM) is 2.9 nM and 4.4 nM, respectively. Hence, perturbation of pAMPK reveals that the region of anoikis resistance is marked by intermediate to low levels of pAMPK, high to intermediate levels of pAkt and intermediate levels of pERK.

Following the perturbation of pAMPK, the system was reset and pERK was perturbed in the same way as above. **Figure 7E** depicts the levels of aCaspase (left Y-axis) and autophagosomes (right Y-axis) when pERK is varied from 0 to 10 nM (X-axis). The steady state value of pERK (4.8 nM) in the matrix-deprived state is marked by the blue-dashed vertical line. The model predicted that from 10 – 1.8 nM of pERK, autophagy outweighs apoptosis and the system is resistant to anoikis. This result is partly corroborated by previous work from our lab, where we have shown that low pERK and high pAMPK levels enable survival (Kumar et al., 2019, bioRxiv). However, when the levels of pERK reduces below 1.8 nM, apoptosis outweighs autophagy and the system succumbs to anoikis. **Figure 7F** depicts the levels of pAkt (left Y-axis) and pAMPK (right Y-axis) when pERK is varied from 0 to 10 nM (X-axis). The levels of pAkt and pAMPK at the crossover (where pERK is 1.8 nM) is 3.8 nM and 8.2 nM, respectively.

Therefore, the above results taken together portrays that a heterogeneity in pERK levels: high/low levels pERK along with high pAMPK enable survival as long as levels of pAkt are maintained low. Additionally, the model predicted a heterogeneity in pAMPK: high/low levels of pAMPK along with low pERK determines the shift from survival to death when levels of pAkt are high. Such high levels of pAkt are obtained at critically low levels of pERK.

### Is there any hierarchy among Akt-AMPK-ERK that may influence the cell fate decision?

The previous section elucidated the effects of Akt-AMPK-ERK on the cell fate decision when each of them are perturbed one at a time. Since each protein affects the other two proteins, we queried whether there is any specific combination of perturbation of Akt-AMPK-ERK which would depict effective crossover of apoptosis over autophagy and give further insights on the region of anoikis resistance. For this experiment, the differential equation of each protein was set to zero and the temporal dynamics of apoptosis (levels of aCaspase) and autophagy (levels of autophagosome) were plotted along with low (2 nM), intermediate (5 nM) and high (8 nM) levels of each protein. The temporal dynamics of two unperturbed proteins are also shown along with the pertubed protein. This experiment is done for a time span of 1440 minutes (or 24 h) in matrix-deprived condition.

Along the lines of the previous experiment, the model predicts that apoptosis (black line) surpasses autophagy (dark-green line) for intermediate and high levels of pAkt (**Figure S10A)** and low levels of pAMPK (**Figure S10B)**. Interestingly, none of the perturbations (low/intermediate/high) of ERK depict a crossover of apoptosis over autophagy (**Figure S10C; right panel**) until 1440 minutes (or 24 h). This is because for all levels of pERK, levels of pAkt and pAMPK are maintained low and high (**Figure S10C**), respectively.

The above results motivated us to enquire if pERK is perturbed along with pAkt or pAMPK, does it enable a crossover of apoptosis over autophagy. For this, the differential equation of two proteins were set to zero and the temporal dynamics of apoptosis (levels of aCaspase) and autophagy (levels of autophagosome) were plotted at low (2 nM), intermediate (5 nM) and high (8 nM) levels of those two proteins. To our surprise, when pERK was perturbed along with pAkt, the model predicted that apoptosis surpased autophagy for intermediate and high levels of the dual protein perturbation (**Figure 8A**, right panel). Simultaneously, we observed a gradual decrease in the levels of pAMPK (green line in **Figure 8A**, left panel) with increase in the levels of pAkt and pERK. When pERK was perturbed along with pAMPK, apoptosis surpased autophagy for low levels of the dual protein perturbation (**Figure 8B**, right panel). Additionally, we observed a gradual decrease in the levels of pAkt (red line in **Figure 8B**, left panel) with increase in the levels of pAMPK and pERK. It is interesting to note that, when pERK is either perturbed along with pAkt or pAMPK (**Figure 8A** and **Figure 8B**), the crossover pattern resembles that of individual perturbation of pAkt (**Figure S10A**) or pAMPK (**Figure S10B**). These results indicate that pAkt and pAMPK dominates over pERK while influencing cell fate decisions.

**Figure 8:**
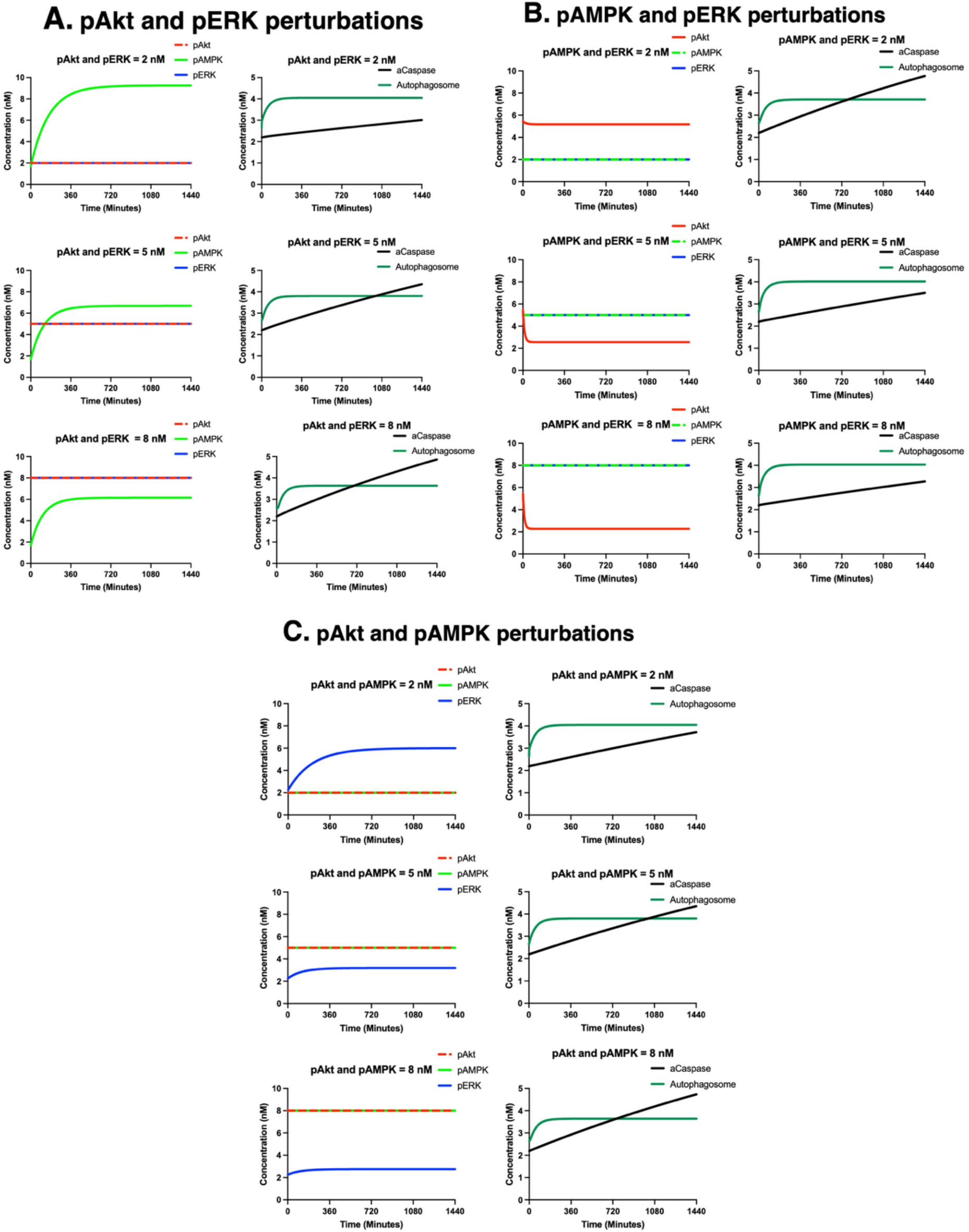
Effect of dual protein perturbation on apoptosis and autophagy. The left panel shows proteins which are maintained in low (2 nM), intermediate (5 nM) and high (8 nM) levels and the right panel shows their effect on apoptosis (aCaspase; black line) and autophagy (autophagosome; dark-green line) for **(A)** pAkt and pERK perturbation, **(B)** pAMPK and pERK perturbation, and **(C)** pAkt and pAMPK perturbation. X axis shows time in minutes. Y axis shows concentration in nM.

The above results further made us curious to question the existance of any hierarchy among pAkt and pAMPK. When dual perturbations of pAkt and pAMPK were simulated, the model predicted crossover of apoptosis over autophagy only for intermediate and high levels of dual perturbations (**Figure 8C**, right panel). It is interesting to note that low levels of dual perturbations of pAkt and pAMPK no longer depicted the crossover, as observed previously when pAMPK was perturbed individually (**Figure S10B**, right panel). Additionally, we also observed the gradual decrease of pERK (blue line in **Figure 8C**, left panel) with the increase in dual perturbation of pAkt and pAMPK. This indicated that pAkt dominates over pAMPK while influencing cell fate decisions.

Hence, the above results illustrate that there exists a hierarchy among Akt-AMPK-ERK and pAkt dominates over pAMPK, which further dominates over pERK while influencing cell fate decision pathways like apoptosis and autophagy.

### What combination of Akt-AMPK-ERK favours pro-apoptotic signals over pro-survival signals?

Intrigued by the findings on single and double perturbations of Akt-AMPK-ERK and their imapct on cell fate pathways, we were curious to know when all three proteins were perturbed at the same time, what combinations of Akt-AMPK-ERK favored apoptosis over autophagy and hence the system would succumb to anoikis. Understanding this will enable us to identify key nodes that can be targeted for therapy.

For this experiment, we sampled five different concentrations (1 nM, 3 nM, 5 nM, 7 nM and 10 nM) of three proteins: pAkt, pAMPK and pERK. This resulted in 5^3^ or 125 different combinations of proteins. For each of these 125 combinations of proteins, the levels of aCaspase and autophagosome were computed and the ratio of aCaspase and autophagosome was evaluated. When the ratio was less than 1 (aCaspase < autophagosome), those combinations of proteins were marked to belong to the pro-survival zone. When the ratio was more than 1 (aCaspase > autophagosome), those combinations of proteins were marked to belong to the pro-apoptotic zone. This experiment was done exclusively for the matrix-deprived state.

Among the 125 combinations of pAkt, pAMPK and pERK, we selected eight extreme combinations for simplicity. **Figure 9A** depicts the eight vertices of the 3D figure denote pAkt^Low^pAMPK^Low^pERK^Low^ ([1,1,1]; **a**), pAkt^Low^pAMPK^High^pERK^Low^ ([1,10,1]; **b**), pAkt^Low^pAMPK^High^pERK^High^ ([1,10,10]; **c**), pAkt^Low^pAMPK^Low^pERK^High^ ([1,1,10]; **d**), pAkt^High^pAMPK^Low^pERK^Low^ ([10,1,1]; **e**), pAkt^High^pAMPK^High^pERK^Low^ ([10,10,1]; **f**), pAkt^High^pAMPK^High^pERK^High^ ([10,10,10]; **g**), and pAkt^High^pAMPK^Low^pERK^High^ ([10,1,10]; **h**). In order to understand, what combinations of pAkt, pAMPK and pERK contribute towards pro-survival or pro-apoptotic cell fate decisions, we plotted the corresponding cell fate decisions in **Figure 9B**. We observe that for pAkt^Low^pAMPK^Low^pERK^Low^, pAkt^Low^pAMPK^High^pERK^Low^, pAkt^Low^pAMPK^High^pERK^High^, and pAkt^Low^pAMPK^Low^pERK^High^, the model predicted pro-survival phenotype. However, for pAkt^High^pAMPK^Low^pERK^Low^, pAkt^High^pAMPK^High^pERK^Low^, pAkt^High^pAMPK^High^pERK^High^, and pAkt^High^pAMPK^Low^pERK^High^ the model predicted pro-apoptotic phenotype. In **Figure 9B**, the 2D plane in blue (known as the ‘decision boundary’) is the plane where ratio of aCaspase and autophagszome is 1. This plane seperates the three-dimensional space into two zones: the zone with all green dots that depict all those combinations of Akt-AMPK-ERK that contribute to pro-survival signals (where the ratio of aCaspase and autophagszome is less than 1) and the zone with red dots that depict all those combinations of Akt-AMPK-ERK that contribute to pro-apoptotic signals (where the ratio of aCaspase and autophagszome is more than 1). It is interesting to note that this decision boundary splits the 3D space into two zones based on the levels pAkt. It depicted that both low/high levels of pAMPK and pERK contributed to pro-survival signals until pAkt levels are maintained below 4 nM. Above 4 nM of pAkt, all combinations of pAMPK and pERK resulted in pro-apoptotic signals. This experiment, corroborating with our previous results, confirmed that the status of pAkt emerged as a deciding factor – increase in pAkt shifts the system from pro-survival to pro-apoptotic state.

**Figure 9:**
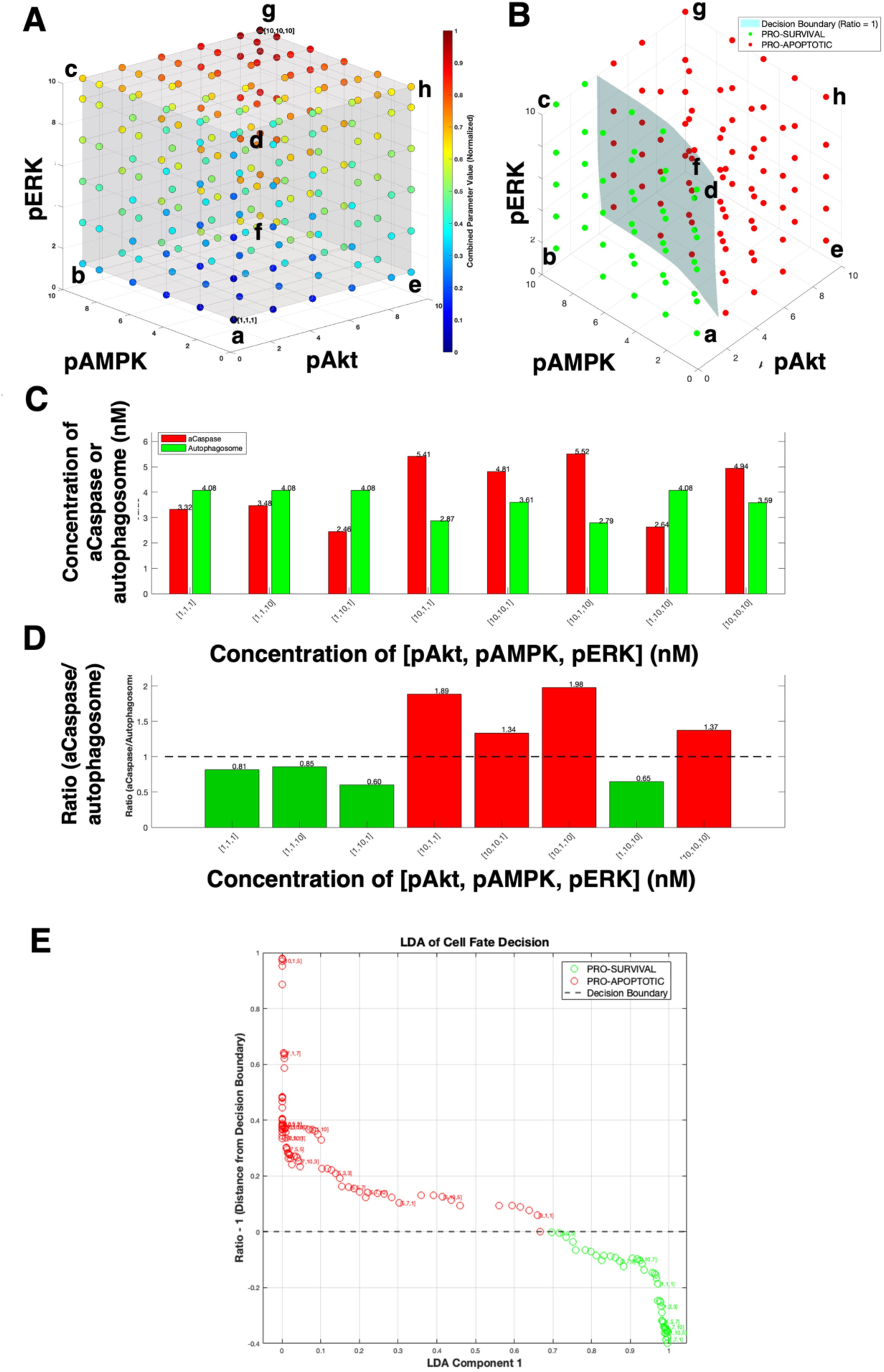
pAkt impacts the balance between apoptosis and autophagy and classifies the cell fate decision into pro-survival and pro-apoptotic zones in the matrix-deprived state. **(A)** A three-dimensional space with 125 different combinations of pAkt (X axis), pAMPK (Y axis) and pERK (Z axis). The coordinates [1,1,1] shows the lowest concentration of pAkt, pAMPK and pERK (colored deep blue) and [10,10,10] shows the highest concentration (colored deep red). The intermediate concentrations, 3 nM, 5 nM and 7 nM, are depicted by intermediate color ranges. The heat map color bar shows a normalized value between 0 and 1. The normalization was done in the following manner: the point [1,1,1] has a combined value of 3, which normalizes to (3-3)/(30-3) = 0, resulting in deep blue color. The point [10,10,10] has a combined value of 30, which normalizes to (30-3)/(30-3) = 1, resulting in deep red. All other points fall somewhere in between on the scale. For example, [5,5,5] has a combined value of 15, normalizing to (15-3)/30-3) = 12/27 ∼ 0.44, resulting in greenish-cyan color. **(B)** For each of these 125 combinations of proteins, the levels of aCaspase and autophagosome were computed and the ratio of aCaspase and autophagosome was evaluated. When the ratio was less than 1 (aCaspase < autophagosome), those combinations of proteins were marked to belong to the pro-survival zone (marked with green dots). When the ratio was more than 1 (aCaspase > autophagosome), those combinations of proteins were marked to belong to the pro-apoptotic zone (marked with red dots). The two-dimensional surface (blue) is called the decision boundary and it marks the plane where the ratio is equal to 1. **(C)** Effects of different concentrations of pAkt, pAMPK and pERK (along the X axis) on aCaspase (red) and autophagosome (green) (along the Y axis). **(D)** Effects of different concentrations of pAkt, pAMPK and pERK (along the X axis) on the ratio of aCaspase/autophagosome (green) (along the Y axis). Ratio below 1 marks the pro-survial zone (green) and ratio above 1 marks the pro-apoptotic zone (red). **(E)** Linear discriminant analysis of 125 combinations of pAkt, pAMPK and pERK and the resultant cell fate outcome. Each circle represents one combination, with colors indicating pro-survival zone (green, ratio < 1) and pro-apoptotic zone (red, ratio > 1). X-axis (LDA Component 1) shows the discriminant score representing the optimal linear combination of pAkt, pAMPK, and pERK that maximally separates cell fate classes. Higher values indicate combinations more favorable to cell survival; lower values indicate combinations more favorable to apoptosis. Y-axis (Ratio - 1, Distance from Decision Boundary) shows distance from the cell fate decision threshold, calculated as the aCaspase/Autophagosome ratio minus 1. Positive values (above dashed line) indicate apoptotic dominance (aCaspase > Autophagosome); negative values (below dashed line) indicate survival dominance (Autophagosome > aCaspase). The horizontal dashed line at y = 0 represents the decision boundary which shows the critical threshold where aCaspase and Autophagosome levels are equal (ratio = 1), separating survival and apoptotic cell fate domains.

In the above results, we found that pAkt emerged as a deciding factor among the three proteins which majorly imapcts the cell fate decision. However, it is not clearly understood what balance of apoptosis and autophagy dictates this decision. Towards understanding this, we computed the levels of aCaspase and autophagosome and plotted them for each of the above eight combinations of proteins. This experiment was also done for the matrix-deprived state and the steady state value for 1440 minutes (or 24 h) were plotted.

**Figure 9C** depicts the levels of aCaspase and autophagosome for all the eight combinations of pAkt, pAMPK and pERK. The labels on the X-axis show the combination of these proteins. The Y-axis shows the levels of aCaspase (red) and autophagosome (green). Following the combinations sequencially in **Figure 9C**, we observed that when the levels of all three proteins was low, levels of autophagosome dominated over levels of aCaspase. When pERK levels were maintained high in addition to low pAkt and pAMPK, a slight increase in levels of aCaspase was noticed, but levels of autophagosome still surpassed levels of aCaspase. When levels of pAMPK were maintained high in addition to low levels of pAkt and pERK, it was interesting to note that there was a decline in the levels aCaspase as compared to the previous two combination, but levels of autophagosome remained unchanged. When levels of pAkt were maintained high, in addition to low levels of pAMPK and pERK, the levels of aCaspase surpassed levels of autophagosome. When both pAkt and pAMPK were maintained high along with low pERK, there was a reduction in aCasapse and increase in levels of autophagosome as compared to the previous case, but apoptosis still dominated over autophagy. When pAkt and pERK were maintained high along with low pAMPK, levels of aCaspase was greater than levels of autophagosome. When pAMPK and pERK was maintained high along with low pAkt, levels of autophagosome was more than aCaspase. When all three levels of proteins were maintained high, levels of aCaspase was more than autophagosome. Previously, we have shown that majority of MDA-MB-231 cells survive upon matrix deprivation and show enhanced autophagy (**Figure 1 and Figure 2**). Hence, we know that autophagy is prominently induced in the system as compared to apoptosis. Hence, when autophagy is high enough, apoptosis is low (as seen in combination 1, 2, 3 and 7; **Figure 9C**), whereas when autophagy is low (as seen in combinations 4,5,6 and 8; **Figure 9C**) apoptosis surpasses autophagy and the model predicts cell death. **Figure 9D** depicts the ratio of aCaspase and autophagosome for all eight combinations of proteins and coroborates the above results.

Our system consists of three key variables (Akt, AMPK and ERK) and each of them affect each other and their combined effect influences cell fate. This enhances the complexity of the above analyses. To minimize the complexity, we attempted methods of dimensionality reduction. Linear discriminant analysis (LDA) and principal component analysis (PCA) are ways of dimensionality reduction. We chose to perform a LDA on cell fate. As compared to PCA which is unsupervised and it does not consider the class difference of survival or death, linear discriminant analysis is supervised, and it preserves the class difference. LDA helps to reduce the dimensionality of three different variables (Akt, AMPK, ERK) by computing a weighted linear combination of all three and then evaluate its effect on cell fate. LDA maximizes between-class variance while minimizing within class variance. LDA analysis depicts that the combination of proteins that indicates towards most apoptotic cells (farthest away and above the decision boundary) comprise of high levels of pAkt, low levels of pAMPK and all concentrations of pERK. Alternately, the combination of proteins that indicates towards most autophagic cells (farthest away and lower the decision boundary) comprise of lowest levels of pAkt, highest levels of pAMPK and all concentrations of pERK (**Figure 9E**). These results corroborate the previous outcomes and ascertain the superiority of Akt on cell fate decisions.

### How does the feedback between Akt-AMPK-ERK impact cell fate decisions?

The previous sections indicate several molecular signatures that favors anoikis. These proteins interact with each other indirectly. There is either a kinase or a phoshatase involved in each case. We hypothesized that perturbing the levels of these kinases and phosphatases may also contribute towards anoikis. Literature shows that PHLPP2 is involved in the negative feedback from AMPK to Akt and PP2Cα is involved in the negative feedback from Akt to AMPK (Saha et al., 2018). PEA15 and TFEB may be involved in the inhibitory feedback from AMPK to ERK and vive versa (Hindupur et al., 2014; Kumar et al., BioRxiv, 2019). Molecular players that are involved in the Akt/ERK crosstalk is not yet known. These intermediate players affect the strength of the feedbacks. In order to study the effect of intermediate players on cell fate decisions, the strength of the feedbacks were altered and their effect on cell fate decisions were studied.

Each of these feedbacks are Hill functions and hence they vary from 0 to 1. **Figure 10Ai** shows that the original strength of the feedback from AMPK to Akt is 0.97 (as written in the bracket). The model is simulated by varying the strength of this feedback from 0.1 to 0.9 at an interval of 0.1, and the dynamics of pAkt, pAMPK, pERK and cell fate were plotted. Since the original strength of this feedback is 0.97, a value of 0.1 indicated least strength of this feedback which is synonymous to most effective knockdown of the intermediate player responsible for that feedback. Varying the strength of this feedback translates to different degrees of knockdown of the intemediate player. By varying the strength of this feedback from 0.1 to 0.9 (**Figure 10Ai**), the model predicted the dynamics the pAkt (**Figure 10Aii**), pAMPK (**Figure 10Aiii**), pERK (**Figure 10Aiv**) and cell fate (**Figure 10Av**). When the feedback was varied from low strength to high strength, the model predicted that the levels of pAkt ranged from 8 nM to 3 nM (**Figure 10Aii**), levels of pAMPK ranged from 7 nM to 8 nM (**Figure 10Aiii**), levels of pERK ranged from 2.5 nM to 4.5 nM (**Figure 10Aiv**). The most effective crossover between the levels of aCaspase and autophagosome happened at the lowest strength of this feedback (**Figure 10Av**). Altering the strength of this feedback primarily affects pAkt. However, the model also shows that different levels of pAkt impacts pAMPK and pERK. This also causes a gradient of cell fate, the most effective one being that of the lowest strength. The same effect was observed when the negative feedback from pERK to pAkt is delinked (**Figure 10B**). However, for no other feedback perturbation (**Figure S11A-D**) , the crossover between apoptosis and autophagy was seen. Hence, the above results depict that in addition to perturbing Akt, the feedbacks affecting Akt can also be targeted to influence cell fate decisions.

**Figure 10:**
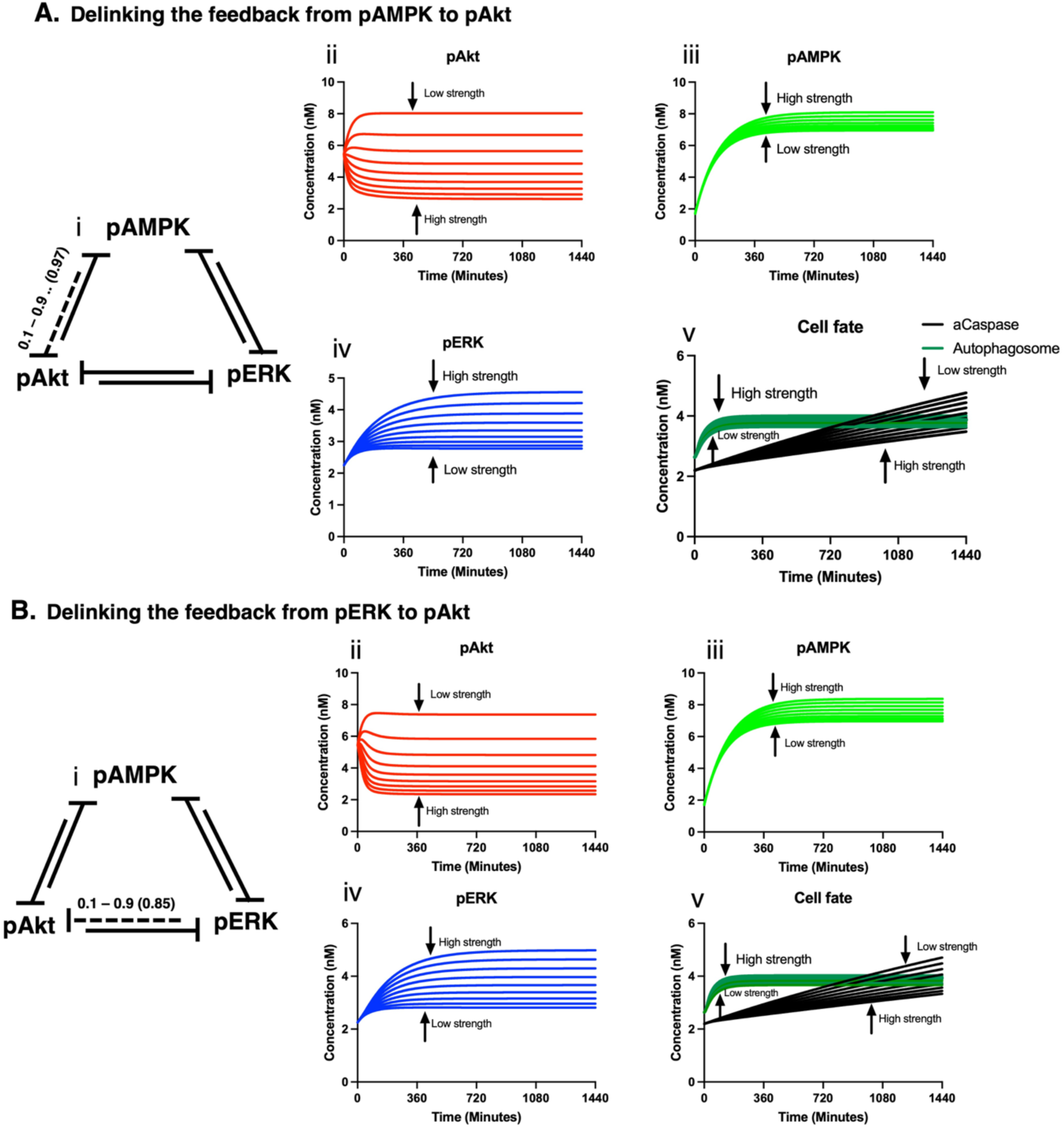
The effect of feedback perturbation on pAkt, pAMPK, pERK and cell fate. (A) **(i)** Delinking the feedback from pAMPK to pAkt**, (ii)** Effect on pAkt, **(iii)** Effect on pAMPK, **(iv)** Effect on pERK and **(v)** Effect on Cell fate. **(B) (i)** Delinking the feedback from pERK to pAkt, **(ii)** Effect on pAkt, **(iii)** Effect on pAMPK, **(iv)** Effect on pERK and **(v)** Effect on Cell fate.

## DISCUSSION

Many cancer cells detach from the extracellular matrix (ECM); however most die due to anoikis and other stresses in the circulation, while some adapt and survive, initiating metastasis. Cancer cells acquire resistance to anoikis by altering several signaling pathways. Understanding these altered pathways that contribute to cell survival is crucial towards combating metastasis. Apoptosis is responsible for cell death, while autophagy generally plays a protective role in cells, albeit excessive autophagic flux or disrupted autophagy leads to cell death. In this study, we see that upon matrix deprivation, anoikis-resistant MDA-MB-231 cells show an enhanced level of autophagy that likely enables circumvention of apoptosis; however, the signaling pathways responsible for this outcome remains elusive. Though some studies have investigated cell fate decisions as a balance between apoptosis and autophagy,^13,14^ this has not been explored extensively in the context of matrix-deprived cancer cells. This study investigates the interplay among Akt, AMPK, and ERK signaling pathways in regulating the balance between apoptosis and autophagy which ultimately affects anoikis resistance.

We began the study by evaluating the status of apoptosis and autophagy in matrix-deprived MDA-MB-231 breast cancer cells. Consistent with our data of AnnexinV/PI and caspase 3/7 activity-based apoptosis assays, literature also shows that MDA-MB-231 cells do not show extensive cell death upon matrix deprivation.^27,28^ However, we observed ∼30% of matrix-deprived MDA-MB-231 show low caspase 3/7 activity. For a long time, executioner-caspase activation has been considered a point-of-no-return in apoptosis. However, recent studies report survival from caspase activation after treatment with drugs or radiation.^29^ One of the very first evidence was shown by Belloc et al. that a slight significant DEVDase activation was also detected in viable cells, though bulk of the DEVDase activity was found in apoptotic cells.^36^ This intrigued us to hypothesize that enhanced autophagy upon matrix deprivation may serve as a barrier for apoptosis and aid in survival. Reviews have documented the paradoxical role of caspase-3 in regulating cell survival.^37^ and promote cytoprotective autophagy during non-lethal stress conditions in human breast cancer cells.^30^ Indeed, our results also depict that there is enhanced autophagy upon matrix deprivation that overcomes low caspase 3/7 activity and aids in cell survival. These results intrigued us to ask how the balance between autophagy and apoptosis is regulated in matrix-deprived cancer cells which enables their survival and subsequent metastasis. Towards understanding this, we formulated a mathematical model of anoikis resistance capturing the effects of matrix deprivation on apoptosis (death) and autophagy (survival) via Akt-AMPK-ERK interplay – 3 key players that couple cell signaling perturbations to cell-fate decisions.

Following model calibration, validation and several mathematical analyses, the model was used to understand what levels of pAkt, pAMPK and pERK can bifurcate the cell fate into survival vs death. Concurrent with previous experimental data^6^, the model predicted that intermediate to high levels of pAkt and low levels of pAMPK trigger cell death. Interestingly, the model predicted a heterogeneity in pERK levels: high/low levels pERK along with high pAMPK enable survival as long as levels of pAkt are maintained low. This result is also partly corroborated by previous work from our lab, where we have shown that low pERK and high pAMPK levels enable survival.^12^ Additionally, the model predicted a heterogeneity in pAMPK: high/low levels of pAMPK along with low pERK determines the shift from survival to death when levels of pAkt are high. Such high levels of pAkt are obtained at critically low levels of pERK. Additionally, we have also shown previously that hyperactivation of Akt is detrimental to matrix-deprived breast cancer cells.^6^ Similarly, Akt hyperactivation has been also shown to induce cell death in chronic lymphocytic leukemia.^38^ Pondering upon these results, we hypothesize that there may exist a threshold of low pERK below which activation of Akt dominates over AMPK to shift the balance in favour of cell death. Low levels of pERK may contribute to cell survival until pAkt is maintained low and pAMPK is high. It may be beyond the scope of experimental techniques to quantify this threshold of low pERK which may contribute towards the activation of Akt and hence the switch from cell survival to cell death. However, this theoretical study indicates towards the possibility of critically low levels of pERK, which leads to high pAkt and favor cell death. The data reveals that matrix-detached cancer cells exhibit complex signaling interactions, where the balance of these interactions dictates cell fate, while cellular heterogeneity arising from these cross-talks might potentially complicate cancer treatment.

Following the above findings, we were curious to know whether the zones of cell fate varied based on a single protein perturbed at a time or more than one protein perturbed simultaneously. This query banked on the fact that the cell fate decisions does not depend only on the perturbed protein but also depends on the effect that the perturbed protein has on the other proteins. Previously, we observed that when the levels of pERK were maintained low, intermediate or high, none of the perturbation depicted the crossover between aCaspase and autophagosome, thereby supporting cell survival (**Figure S10C**). However, when pERK was perturbed either with pAkt or pAMPK (**Figure 8A-B**), we observed the cell fate patterns similar to independent perturbations of pAkt and pAMPK (**Figure S10A-B**). This indicated towards the existence of a hierarchy among Akt-AMPK-ERK while deciding the cell fate. Our results from the dual perturbation studies indicated that pAkt dominated over pAMPK which further dominated over pERK while contributing to cell fate decisions. When all the three proteins are pertubed at the same time, we acquired a three-dimensional space defining the cell fate. In this case too, we found that pAkt split this three-dimensional space into two zones of pro-survival and pro-apoptotic (**Figure 9A-B)**. Additionally, feedback perturbation study also revealed that removing inhibitory feedbacks from AMPK to Akt and ERK to Akt, predicted cell death (**Figure 10A-B**). Thus, in this study, Akt emerged as the deciding factor in influencing cell fate pathways like apoptosis and autophagy in matrix-deprived cancer cells. These results suggest the possibility of narrowing down to key molecular targets to attack matrix-deprived cancer cells.

Lastly, we wanted to understand the balance between autophagy and apoptosis in determining anoikis-resistance. By sampling eight extreme combinations of pAkt, pAMPK and pERK (**Figure 9C**), we noticed that as long as autophagy was high, it resisted cell death, and apoptosis is low. But as autophagy decreased, it led to increase in apoptosis and the model predicted cell death. A dynamical model with quantitative measurements of autophagy and apoptosis in rat kidney proximal tubular cells modelled apoptosis as an all-or-none commitment of individual cells indicating that cells shift between survival and death by switching on or off apoptosis. However, autophagy was mainly incorporated to buffer stress and alter the apoptotic threshold or the point of cell death.^14^ Furthermore, reviews on “cellular decisions between apoptosis and autophagy” described autophagy as a homeostasis, cytoprotective process which attempts at suppressing apoptosis and delaying death.^39^ Since MDA-MB-231 cells are inherently anoikis resistant, cell death pathways like apoptosis may be supressed by autophagy and this may majorly contribute to cell survival. An attempt to inhibit autophagy may enhance cell death. Indeed, using human colorectal carcinoma cell line (HCT116) Fitzwalter et al. depict that autophagy inhibitors can improve anticancer drugs by increasing sensitivity to apoptosis.^40^

### Limitations of the study

Though there are several interesting insights in this work, there are a few limitations too. In this study we did not consider the effect of aCaspases or autophagosomes on pAkt, pAMPK and pERK. Here, only the proteins influence the cell fate markers. Additionally, we did not look into the crosstalk between autophagy and apoptosis as described previously.^39,41^ Here, we described the balance between autophagy and apoptosis as a cell fate for cell survival or cell death. Although the trend of these cross-talks appears to hold true in several cancer lines across various contexts, we have trained the model primarily with Western blotting data from MDA-MB-231 cell line subjected to matrix deprivation for 24 h to avoid inconsistencies caused by varying genetic background, time points, contexts and antibodies. Nevertheless, the model is designed to be flexible; it can be updated with independent datasets and generalized to fit different problem statements. Though, it is unlikely that such predictive systems would be applicable to all cells, as stated previously by other similar work.^42^ Additionally, both time and status of matrix are independent variables, and the proteins and cell fate markers depend on the time and status of matrix. However, the transition from attached state to matrix-deprived state is also a time-dependent phenomena. It will be interesting to introduce matrix as a time dependent variable and then check the effect of matrix on the system. Lastly, due to the availability of experimental data at limited timepoints, parameter estimation was performed manually and the model fitting to experimental data was done qualitatively. It will be very helpful to design future experiments which would be conducive for mathematical modeling work and hence aid in the understanding a biological phenomenon jointly.

## RESOURCE AVAILABILITY

### Lead contact

Requests for further information and resources should be directed to and will be fulfilled by the lead contact, Anu Rangarajan.

### Materials availability

This study did not generate new unique reagents.

### Data and code availability

The western blot data used in this paper will be shared by the lead contact upon request.

All original code is available in this paper’s supplemental information.

Any additional information required to reanalyze the data reported in this paper is available from the lead contact upon request.

## ACKNOWLEDGMENTS

We acknowledge Manipa Saha and Saurav Kumar for experimental data that was used to build this mathematical model. We acknowledge the flow cytometry facility of DBG department as well as division of biological sciences. We also acknowledge the microscopy facility of the division of biological science. We acknowledge Diptanshu Banerjee for helping with experiments. S.M. acknowledges the Prime Minister Research Fellowship (PMRF). Funds from Wellcome Trust DBT India Alliance Senior Research Fellowship (500112-Z-09-Z to AR) and funds from the DBT-IISc partnership programme of the Department of Biotechnology, Govt of India, for experimental data used in this manuscript are acknowledged. Infrastructural support from the UGC and DST-FIST to the Department of DBG is also acknowledged.

## AUTHOR CONTRIBUTIONS

S.M., and A.R. conceptualized and designed the study. S.M. performed all the experiments, constructed the model in MATLAB, performed all the computer simulations, assembled and analyzed the data and scripted the original draft of the manuscript. A.C. supervised the model building and all the mathematical analyses for model construction. S.M., A.C, M.K.J, N.C., and A.R. were involved in interpreting the results and critical discussion. A.R., M.K.J, N.C. edited and revised the manuscript.

## DECLARATION OF GENERATIVE AI AND AI-ASSISTED TECHNOLOGIES

No generative AI or AI-assisted technologies were used during the preparation of this work.

## SUPPLEMENTAL INFORMATION

**Supplementary file. Figures S1–S11, Tables S1–S4, Model equations, MATLAB Code and supplemental references**

## MATERIALS AND METHODS

KEY RESOURCES TABLE.

KEY RESOURCES TABLE
| REAGENT OR RESOURCE | SOURCE | IDENTIFIER |
| --- | --- | --- |
| **Antibodies** |  |  |
| Rabbit polyclonal phospho-p44/42 Erk1/2 antibody | Cell Signaling Technology | CST#9101; RRID:AB_331646 |
| Rabbit monoclonal p44/42 Erk1/2 antibody | Cell Signaling Technology | CST#4695; RRID:AB_390779 |
| Rabbit monoclonal phospho-Akt (Ser473) antibody | Cell Signaling Technology | CST#4060; RRID:AB_2315049 |
| Rabbit polyclonal Akt antibody | Cell Signaling Technology | CST#9272; RRID:AB_329827 |
| Mouse monoclonal anti-α-tubulin antibody | Sigma-Aldrich | CP06; RRID:AB_2617116 |
| Goat polyclonal secondary Anti-Rabbit IgG (H+L)-HRP Conjugate | Jackson ImmunoResearch Laboratories | 111-035-045; RRID:AB_2337938 |
| Goat polyclonal secondary Anti-Mouse IgG (H+L)-HRP Conjugate | Jackson ImmunoResearch Laboratories | 115-035-062; RRID:AB_2338504 |
| **Bacterial and virus strains** |  |  |
| - |  |  |
| **Biological samples** |  |  |
| - |  |  |
| **Chemicals, peptides, and recombinant proteins** |  |  |
| MEK/ERK inhibitor, PD98059 | Sigma-Aldrich | 513000 |
| Dimethyl sulfoxide (DMSO) | Sigma-Aldrich | 317257 |
| Noble Agar | BD Difco | 214230 |
| Hydrogen peroxide (H<sub>2</sub>O<sub>2</sub>) | Merck EMPLURA | 1.93407 |

| Critical commercial assays |  |  |
| --- | --- | --- |
| Autophagy assay kit | Sigma Aldrich | MAK138 |
| CellEvent Caspase-3/7 detection kit | Invitrogen | C10431 |
| Dead cell apoptosis kit with Annexin V Alexa Fluor 488 and propidium iodide | Invitrogen | V13245 |
| Deposited data |  |  |
| - |  |  |
| Experimental models: Cell lines |  |  |
| MDA-MB-231 | ATCC | HTB-26<br>RRID:CVCL_0062 |
| Experimental models: Organisms and strains |  |  |
| - |  |  |
| Oligonucleotides |  |  |
| - |  |  |
| Recombinant DNA |  |  |
| - |  |  |
| Software and algorithms |  |  |
| MATLAB R2023a software | Mathworks | http://www.mathworks.com/products/matlab/<br>MATLAB;<br>RRID:SCR_001622 |
| GraphPad Prism 10 software | GraphPad | https://www.graphpad.com/<br>GraphPad Prism;<br>RRID:SCR_002798 |
| Fiji/ImageJ | ImageJ | http://fiji.sc<br>Fiji;<br>RRID:SCR_002285 |
| Other |  |  |
| - |  |  |

## EXPERIMENTAL MODEL AND STUDY PARTICIPANT DETAILS

### METHOD DETAILS

#### Extraction of experimental data

*In vitro* experimental data retrieved from previous work of our laboratory (Saha et al., 2018; Kumar et al., BioRxiv, 2019) as well as published literature (Stulpinas et al., 2023) were used for model calibration and validation. Some additional experiments (**Figure 1-2**) were also performed to build the model. All the data used for model calibration and validation are summarized in the **Table S1.** In brief, the data used for model calibration comprise of the effect of matrix deprivation on pAkt, pAMPK, pERK, autophagy and apoptosis, across multiple timepoints. The data used for model validation consist of the effect of Akt hyperactivation and AMPK inhibition on apoptosis and autophagy. Additionally, the data also consists of the effect of PHLPP2 knockdown on pAkt, pAMPK (in the presence or absence of Akt inhibition), apoptosis and autophagy (was also used for validation).

#### Formulation of anoikis resistance mathematical model

All the variables of the anoikis resistance model are listed in **Table S2.** The generic deterministic equations representing the temporal dynamics of pAkt, pAMPK, pERK, apoptosis and autophagy are given by:

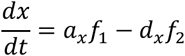

where 𝑎_x_ 𝑎𝑛𝑑 𝑑_x_ represent the basal activation and deactivation rates of the component 𝑥 and 𝑓_1_𝑎𝑛𝑑 𝑓_2_ are two functions representing the regulation of 𝑥’s activation and deactivation due to crosstalk between 𝑥 and other components. The ordinary differential equations (ODEs) were formulated based on kinetic rate laws and mass balance of signaling proteins. Hill functions were used to capture important feedback and crosstalks in the network. Details on the model equations are elaborated in the **Supplemental Information section** under ‘**Model equations’**. Dynamic solutions of these ODEs were obtained using ODE15s solver in MATLAB R2023a software. The initial conditions of the model and the steady state values of matrix-attached and matrix-deprived condition are listed in **Table S3.**

#### Parameter estimation

The parameters of this mathematical model were either retrieved from literature or estimated sequentially through trial and error by fitting the model within the range defined by experimental data (as shown in **Table S1A**). Since all the experimental data were obtained by western blotting, it was challenging to retrieve the data at regularly-spaced time intervals. Hence, most of the experimental data used for this study belong to early time points like 10 mins or 1 h or 2 h and later time points like 8 h or 24 h. Since uniformly spaced time points are not available, we performing a qualitative model fitting to the experimental data. To determine the baseline parameter set, we performed a grid search (within a specific range) for parameters associated with hill functions. Hill functions capture the interactions between Akt, AMPK, ERK and their impact on autophagy and apoptosis. These parameters included V (fold change of hill function; 1 - 100), k (threshold of hill function; 0 - 10) and n (hill coefficient; 2 - 8). Additionally, five other parameters namely, k_a_Cas_Basal_ (basal rate of activation of caspases), k_a_Cas (Rate of activation of caspases), k_fpfBasal_(Basal rate of formation of phagophore), k_fpg_(Rate of formation of phagophore) and k_fAgBasal_ (Basal rate of formation of autophagosome) were modified from values available in the literature (Liu et al., 2017). All the parameters are tabulated in **Table S4.**

Model calibration was done by estimating the above parameters to obtain fold change values of pAkt, pAMPK, pERK, autophagy and apoptosis for different conditions (attached vs suspension and treatment vs. control) within the range of experimental data as summarized in **Table S1A.**

The calibrated model was further validated using independent set of experimental data that was not used previously for calibration (as summarized in **Table S1B**).

#### Temporal dynamics of the model

To identify the steady states of the above calibrated and validated model, the model was simulated for 1440 min (24 h) as well as for longer durations (20,000 h). The timepoint of 24 h was chosen because most of the experimental data were available for 24 h timepoint. To understand the dependence of these steady states on the initial conditions, 1000 random initial conditions of pAkt, pAMPK and pERK were sampled from a uniform distribution of range 1-10 nM and were plotted against the steady state values.

#### 3D-Nullcline analysis

A nullcline is a curve on which the derivative of one variable is set to zero. The intersection of nullclines depicts the steady states of the system. To confirm the steady states of the system, 3D-nullclines for pAkt, pAMPK and pERK were obtained by plotting 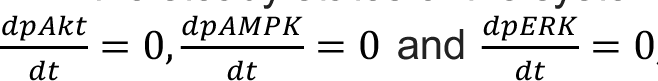 and inline], in a three-dimensional space with axes pAkt (X-axis), pAMPK (Y-axis) and pERK (Z-axis). Since each variable depends on two other variables, each curve is a two-dimensional surface in three-dimensional space. For example, to plot the nullcline of pAkt, 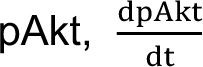 was set to zero, and pAkt was written in term of pAMPK and pERK 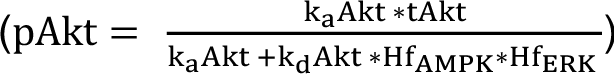. Then, for all possible values of pAMPK and pERK (ranging from 0 to 10), values of pAkt were computed using MATLAB. The resultant 2D surface was plotted in red for matrix-attached condition and for matrix-deprived condition. Likewise, the nullclines for pAMPK and pERK were also plotted separately.

#### Stability analysis

A standard method to study the stability of steady states is linear stability analysis which is done by computing the Jacobian of differential equations and finding the corresponding eigenvalues. However, for a system of complex coupled-ODEs like in this anoikis resistance model, it is challenging to perform a linear stability analysis. Hence, to study the stability of the steady states of this system, steady state perturbation analysis and phase portraits were plotted.

*Steady state perturbation analysis* was performed by perturbing the steady state value (decreasing by 50% or increasing by 50%), and the temporal dynamics for pAkt, pAMPK and pERK in both matrix-attached and matrix-deprived state were plotted against time.

*Phase portraits* consist of vector fields at all points on the phase planes. The vector field at a particular point is computed by evaluating the sign of the derivative of all the variables which are denoted as individual vectors (plotted along the axes of the phase planes). The resultant of all the vectors at that point shows the direction in which the system may evolve if the system began from that point.

#### Flux map analysis

To visualize the relative operational strengths of the feedback and crosstalks under matrix-attached and matrix-deprived conditions, the absolute values of the corresponding Hill functions were evaluated and plotted in the flux maps.

#### Parameter sensitivity analysis

To assess how small perturbation in individual model parameter affect the model variables, parameter sensitivity analysis was done. The sensitivity of a steady-state output signal S to changes in the parameter p can be determined by the following formula:

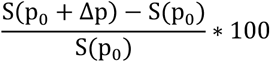

S(p_0_) corresponds to the steady state concentration of the signal S obtained using the initial parameter value p_0_. The parameter of interest p is decreased by 10% (Δ𝑝 = − 0.1 ∗ 𝑝_0_), and the new steady state concentration 𝑆(𝑝_0_ + Δ𝑝) is obtained.

#### Differential activity of proteins

To study the effect of differential activity of Akt-AMPK-ERK on each other and autophagy/apoptosis, the differential equation of each of the proteins was set to zero and the levels of these proteins were varied from 0 to 10 nM. The effects of the differential levels of these proteins were studied on the model outputs.

#### Node and feedback perturbation

To study the effect of single versus double node perturbation, each molecular player (Akt-AMPK-ERK) was set at three different levels (low-intermediate-high), and their effects were studied on the model outputs. Subsequently, the feedback strengths ranging between 0 – 1 were also varied and their effects were studied on the model outputs.

#### Linear discriminant analysis (LDA)

To study how Akt-AMPK-ERK influence cell fate decisions, LDA was performed. This is a supervised dimensionality reduction technique that uses class labels to find the most discriminative directions in the data. LDA seeks to maximize the separation between classes while minimizing within-class variance.

#### Experimental design

##### Cell culture

Human breast cancer cell line MDA-MB-231 (ATCC) was cultured in DMEM (Dulbecco’s Modified Eagle Medium; Sigma-Aldrich) supplemented with 10% FBS (fetal bovine serum; Gibco) containing penicillin (HiMedia) and streptomycin (HiMedia), at 37°C and 5% CO_2_. For matrix-detached (suspension) cultures of 24 hours, cancer cells were seeded on dishes coated with 2% noble agar (Sigma-Aldrich) (Warrier et al., 2023).

##### Pharmacological compounds

Pharmacological compound used in cell culture was MEK/ERK inhibitor, PD98059 (10 μmol/L; referred to as PD in figures). Dimethyl sulfoxide (DMSO) was used as vehicle control for the pharmacological compound. For experiments carried out in matrix-deprived (suspension) condition, pharmacological compound was added during initial seeding. No prior treatment was given.

##### Immunoblotting

For immunoblotting, 5 x 10^5^ cells were seeded in 35 mm dishes coated with noble agar and exposed to matrix deprivation for 24 h. Whole-cell lysates of suspension cells were prepared using RIPA lysis buffer as described before (Hindupur et al., 2014). Suspension cells were collected into a microcentrifuge tubes, spun down to remove media and then lysed using the lysis buffer. Protein concentration was estimated using Bradford method and equal quantity of protein (30 – 50 μg) per lane was resolved by SDS-PAGE after boiling with sample buffer for 2 minutes at 95 °C. Proteins were transferred to PVDF membrane and probed with appropriate antibodies. The membrane was incubated overnight with primary antibody at 4 °C followed by TBST washes and incubated for 2 h with HRP-conjugated secondary antibodies at room temperature. Chemiluminescence (using ECL substrate from Bio-Rad) was used to visualize proteins bands. α-tubulin served as loading control for each run. Representative immunoblots show data consistent with minimally three independent experiments. Densitometric analyses of Western blots were performed using Fiji/ImageJ software. Relative protein levels were quantified by normalizing to loading control. Primary antibodies used in the study were against pERK1/2, tERK, pAkt^S473^, tAkt (Cell Signaling Technology) and a-tubulin (CP06; Merck). HRP-conjugated anti-mouse and anti-rabbit antibodies were obtained from Jackson ImmunoResearch Laboratories.

##### Autophagy assay

Autophagy was measured by autophagy assay kit (MAK138; Sigma Aldrich) according to the manufacturer’s instructions. Briefly, 1.5 x 10^5^ MDA-MB-231 cells were seeded in a 24-well plate in attached and suspended condition. Following 24 h of suspension, cells were stained with 0.4 μL of autophagosome detection reagent in 200 μL of stain buffer and incubated for 30 minutes at 37°C and 5% CO_2_. Cells incubated with stain buffer only were used as negative control. Cells that were exposed to serum-free media for 24 h to induce autophagy and incubated with autophagosome detection reagent were used as positive control. Cells were washed thrice using wash buffer. Both attached and suspended cells were trypsinized before resuspending in PBS for fluorescence-activated cell sorting (FACS). Green (500 – 560 nm) fluorescence emission was measured after illumination with ultra-violet (360 nm) excitation light. Autophagosome-positive cells were sorted and put on a glass slide and covered with cover slip and analyzed using Confocal microscopy. Data were analyzed using Fiji/ImageJ software.

##### Caspase 3/7 activity assay

Caspase 3/7 activity was measured by using CellEvent Caspase-3/7 detection kit (C10431; Invitrogen) according to the manufacturer’s instructions. Briefly, 1.5 x 10^5^ MDA-MB-231 cells were seeded in a 24-well plate in attached and suspended condition. Following 24 h of suspension, cells were rinsed in PBS and stained with 2 μL of caspase 3/7 detection reagent in 200 μL of complete media and incubated for 30 minutes at 37°C and 5% CO_2_. Cells incubated with complete media only were used as negative control. Cells that were exposed to 0.5 mM of H_2_O_2_ for 1 h to induce apoptosis and incubated with caspase 3/7 detection reagent were used as positive control. Cells were washed thrice using PBS. Both attached and suspended cells were trypsinized before resuspending in PBS for fluorescence-activated cell sorting (FACS). PE-Texas Red (610 nm) fluorescence emission was measured after illumination with yellow-green (561 nm) excitation light. Caspase 3/7 positive cells were sorted and put on a glass slide and covered with cover slip and analyzed using Confocal microscopy. Data were analyzed using Fiji/ImageJ software. For dual-staining of caspase 3/7 and autophagosome, the above methods were carried out simultaneously, sorted using FACS and analyzed using confocal microscopy.

##### Apoptosis assay

Apoptosis was measured using dead cell apoptosis kit with Annexin V Alexa Fluor 488 and propidium iodide (V13245; Invitrogen) according to the manufacturer’s instructions. Briefly, 1.5 x 10^5^ MDA-MB-231 cells were seeded in a 12-well plate in attached and suspended condition. Following 24 h of suspension, cells were harvested and washed twice with cold phosphate buffer saline (PBS) and then resuspended in 100 μL of 1X annexin-binding buffer. Following this, 5 μL of Annexin V and 1 uL of 100 μg/mL PI working solution was added to each 100 μL of cell suspension and cells were incubated at room temperature for 15 minutes in the dark. After the incubation period, 400 μL of 1X annexin-binding buffer was added and the samples were gently mixed and kept in ice. Samples were analyzed by flowcytometry. The percentage of live cells was detected by Annexin V(-)/PI(-), necrotic cells were detected by Annexin V(-)/PI(+), early apoptotic cells were detected by Annexin V(+)/PI(-) and late apoptotic cells were detected by Annexin V(+)/PI(+).

## QUANTIFICATION AND STATISTICAL ANALYSIS

Statistical analyses were performed using MATLAB R2023a software using Student’s t test. All data are presented as mean±SEM, where *, P < 0.05; **, P < 0.01; ***, P < 0.001; ns (non-significant), P>0.05.

