## Supplementary file for "Balance between autophagy and apoptosis determines anoikis resistance: a mathematical model on cell fate"

### Supplemental Information

#### Figure S1

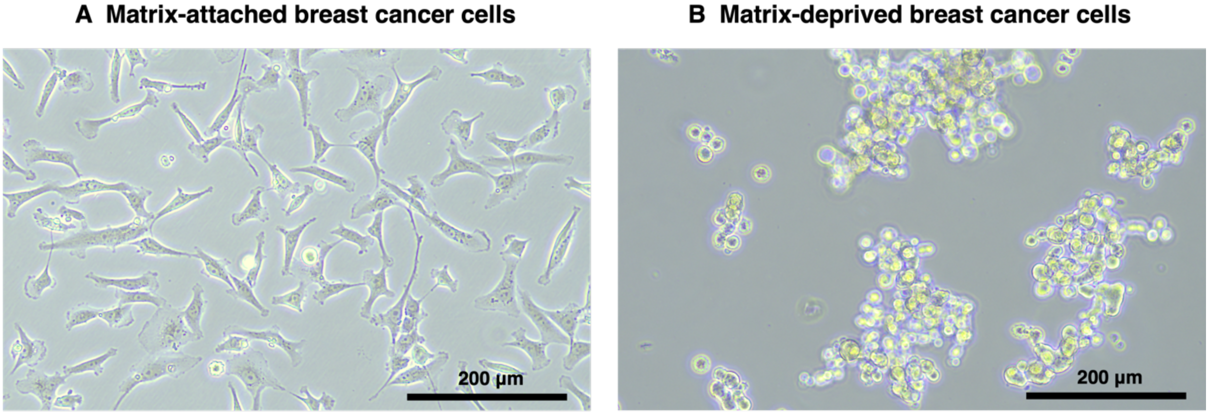


**Figure S1:** Experimental system of study consists of MDA-MB-231 breast cancer cells grown in **(A)** matrix-attached and **(B)** matrix-deprived conditions. Cells are grown on regular tissue culture plastic dishes for matrix-attached condition. Cells are cultured on 2% noble-agar coated dishes for matrix-deprived condition. The figure shows phase contrast images of MDA-MB-231 cells at 10x magnification in matrix-attached and matrix-deprived conditions. Scale bar: 200 μM.

#### Figure S2

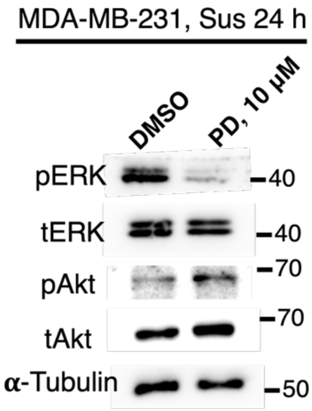


**Figure S2:** ERK inhibition by PD98059 increases pAkt levels in suspension.

MDA-MB-231 cells were subjected to suspension culture (Sus) for 24 h in the presence of vehicle control (DMSO) or 10 μM of MEK/ERK inhibitor (PD98059), n = 2.

#### Figure S3

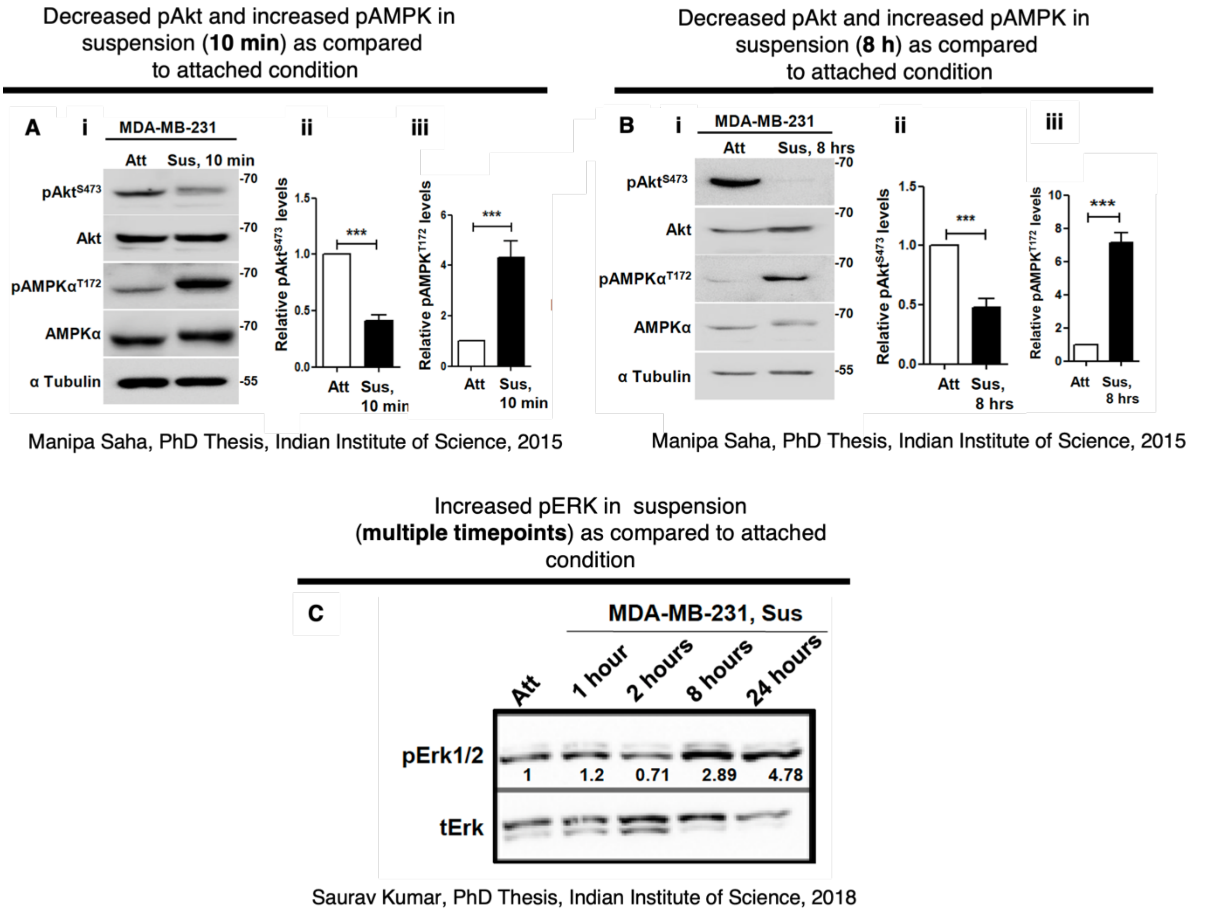


**Figure S3:** Experimental data used for calibration. Effect of matrix deprivation on pAkt, pAMPK, and pERK. Upon matrix deprivation there was decreased pAkt and increased pAMPK when MDA-MB-231 cells were exposed to suspension for (A) 10 mins and (B) 8 h (A-B; reprinted from Saha et al., with permission from AACR). (C) Upon matrix deprivation there was increased levels of pERK.

#### Figure S4

AMPK activation (by genetic techniques) in attached cells decreases pAkt levels

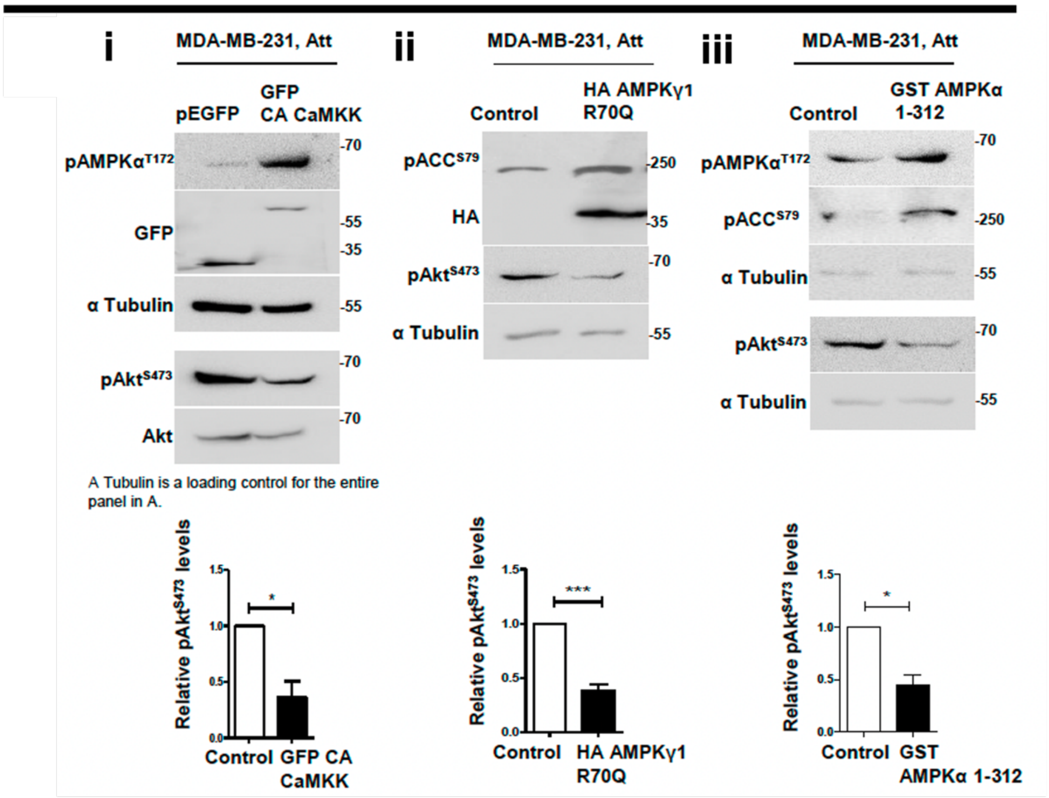


Manipa Saha, PhD Thesis, Indian Institute of Science, 2015

**Figure S4:** Experimental data used for calibration. AMPK activation by genetic techniques in attached cells decreases pAkt levels. (i; reprinted from Saha et al., with permission from AACR).

#### Figure S5

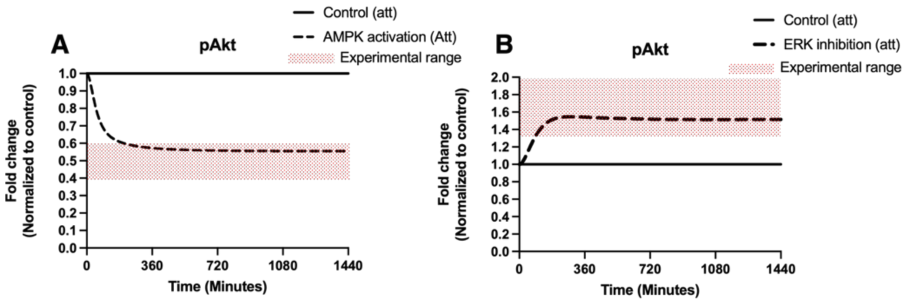


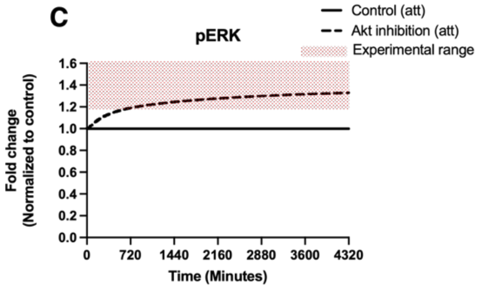


##### Summary of calibration data

| Sr. No. | Variables | Treatment | Condition | Effect |
|---|---|---|---|---|
| 1. | pAkt | Effect of AMPK activation | Att | Decrease |
| 2. | pAkt | Effect of ERK inhibition | Att | Increase |
| 3. | pERK | Effect of Akt inhibition | Att | Increase |

**Figure S5. Model calibration:** The calibration set consists of temporal dynamics of simulated concentration of following protein species in MDA-MB-231 cell line for 24 h period (A) effect of AMPK activation on pAkt in matrix-attached state, (B) effect of ERK inhibition on pAkt in matrix-attached state, (C) effect of Akt inhibition on pERK in matrix-attached state. The black line (solid: control, dashed: perturbed state) depicts the dynamics predicted by the model. X axis depicts time in minutes, Y axis depicts the fold change normalized to control.

#### Figure S6

PHLPP2 knockdown leads to decreased AMPK activity in matrix-detached cells via Akt activation

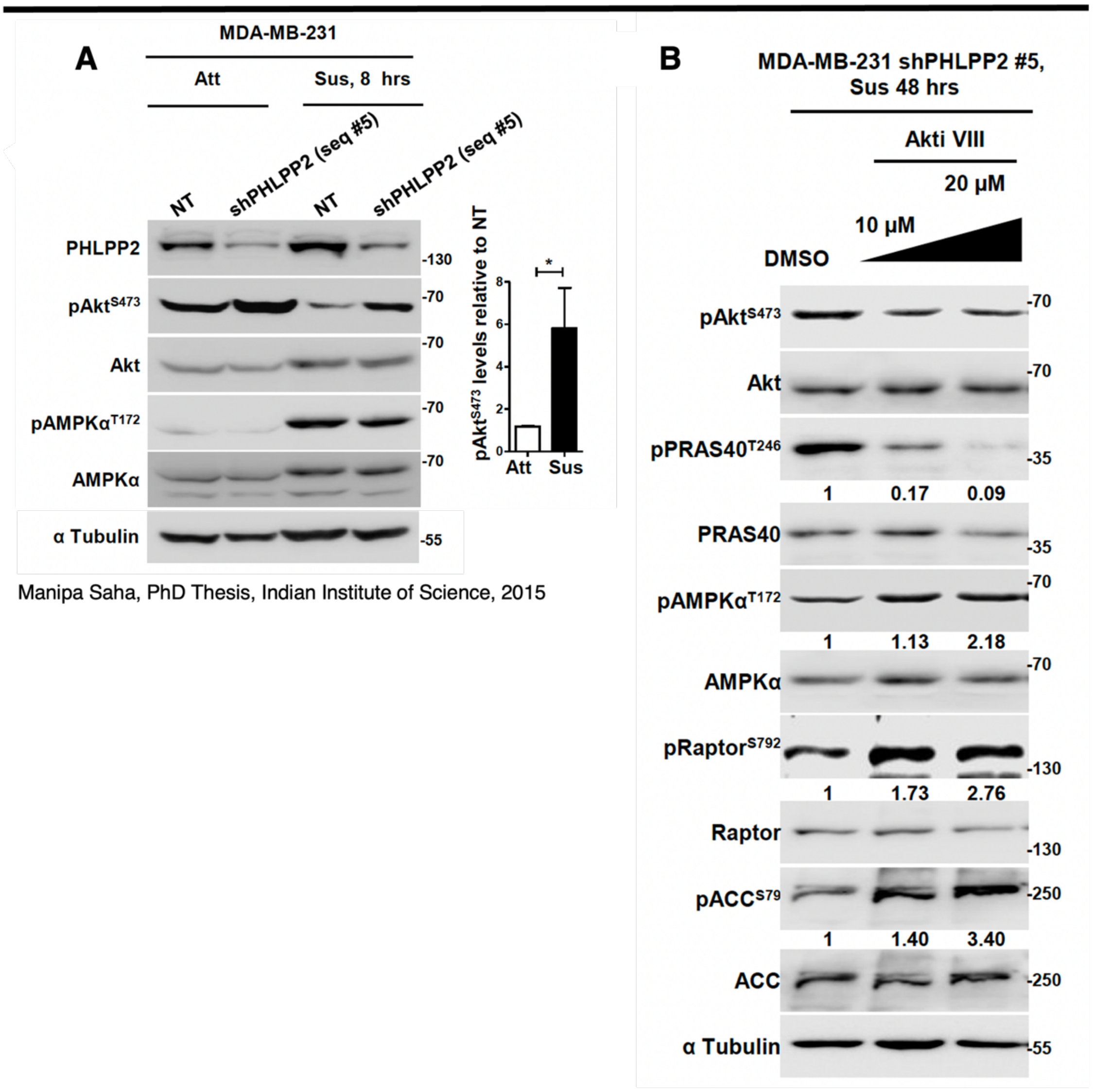


Manipa Saha, PhD Thesis, Indian Institute of Science, 2015

Manipa Saha, PhD Thesis, Indian Institute of Science, 2015

**Figure S6.** Experimental data used for validation. Effect of PHLPP2 KD on pAMPK and pAkt in the (A) absence of Akt inhibitor and (B) presence of Akt inhibitor.

### Figure S7

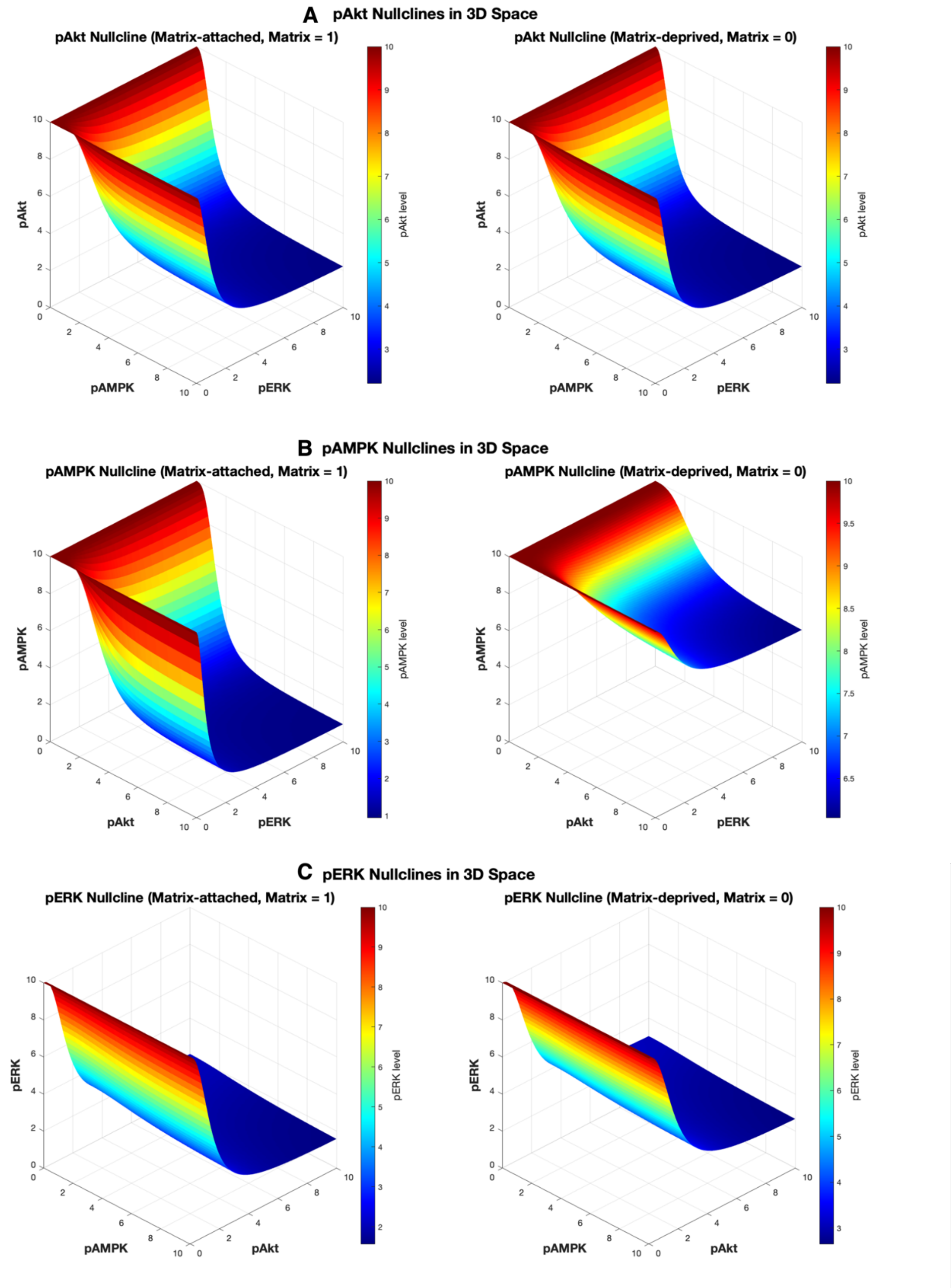


**Figure S7: (A)** 3D-nullcline of pAkt, (B) 3D-nullcline of pAMPK and (C) 3D-nullcline of pERK in matrix-attached (left panel) and matrix-detached (right panel) condition. X axis depicts concentration of pAkt,

#### Figure S8

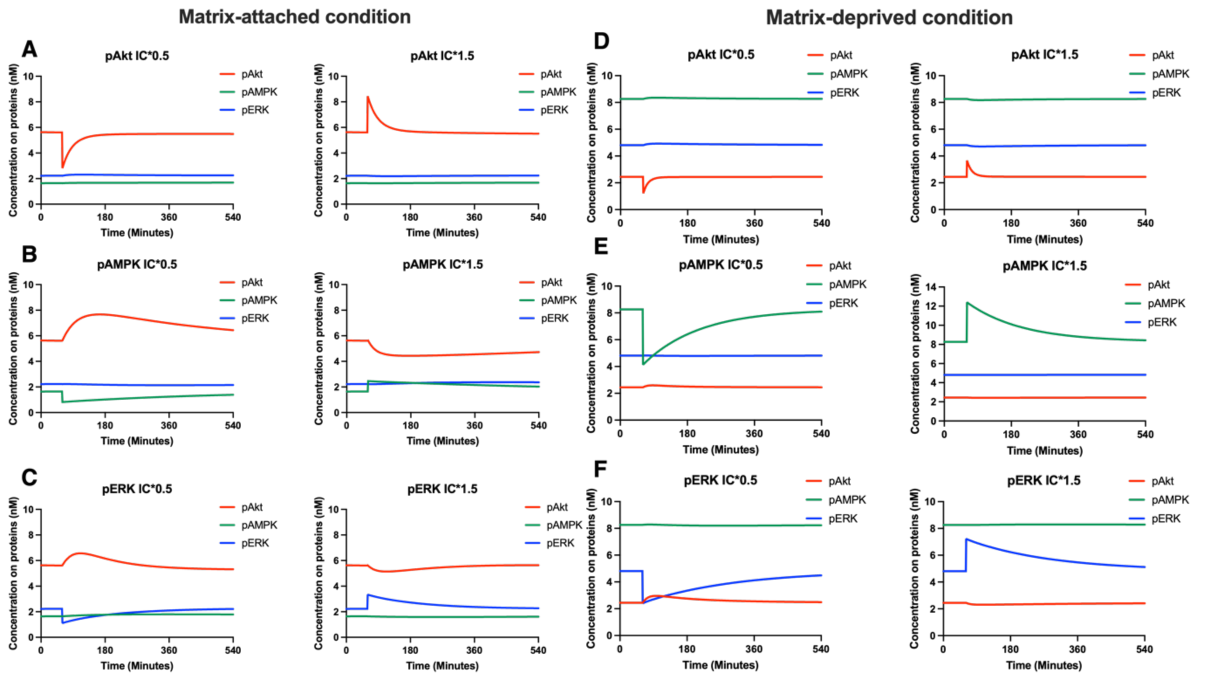


**Figure S8:** Steady state perturbation analysis. The steady states of each protein are decreased or increased by 50% and dynamics of the steady states is studied for matrix-attached (left panel and matrix-deprived (right panel). (A) and (D) correspond to pAkt. (B) and (E) correspond to pAMPK. (C) and (F) correspond to pERK.

#### Figure S9

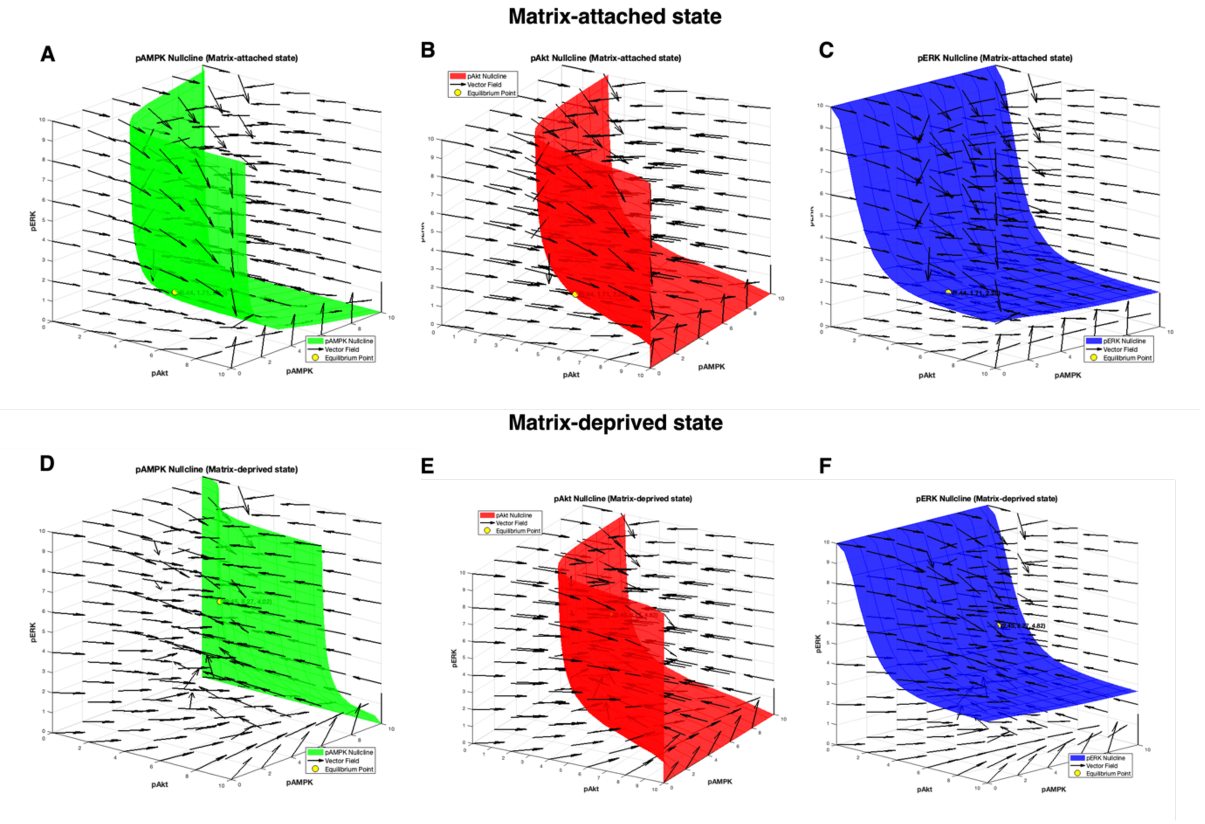

**Figure S9:** Phase portraits of pAMPK (A and D), pAkt (B and E) and pERK (C and F) for matrix-attached (top panel) and matrix-deprived state (bottom panel). Each arrow depicts the vector field at that point.

### Figure S10

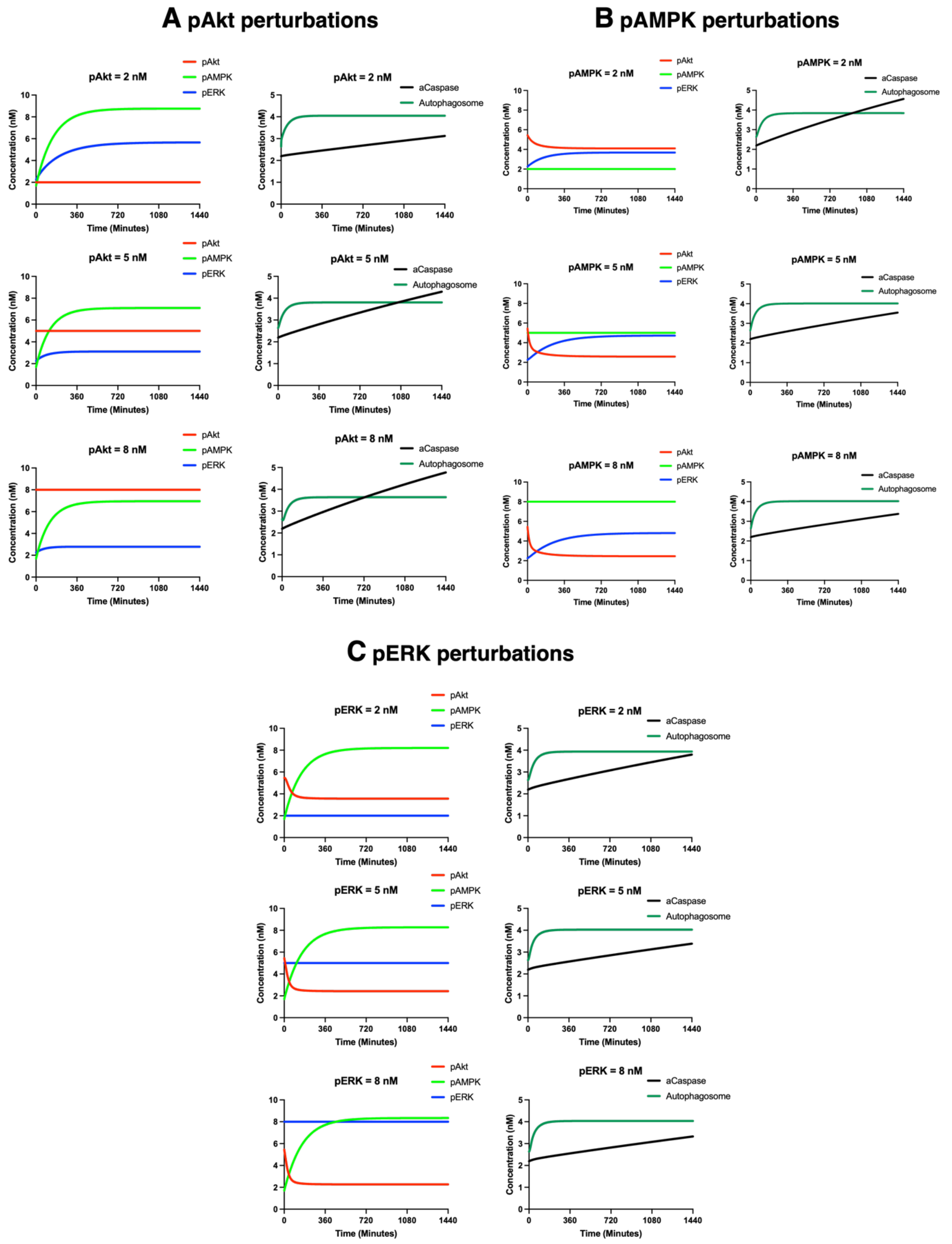


**Figure S10: Effect of single protein perturbation on apoptosis and autophagy.** The left panel shows proteins which are maintained in low (2 nM), intermediate (5 nM) and high (8 nM) levels and the right panel shows their effect on apoptosis (aCaspase; black line) and autophagy (autophagosome; darkgreen line) for **(A)** pAkt perturbation, **(B)** pAMPK perturbation, and **(C)** pERK perturbation. X axis shows time in minutes. Y axis shows concentration in nM.

#### Figure S11

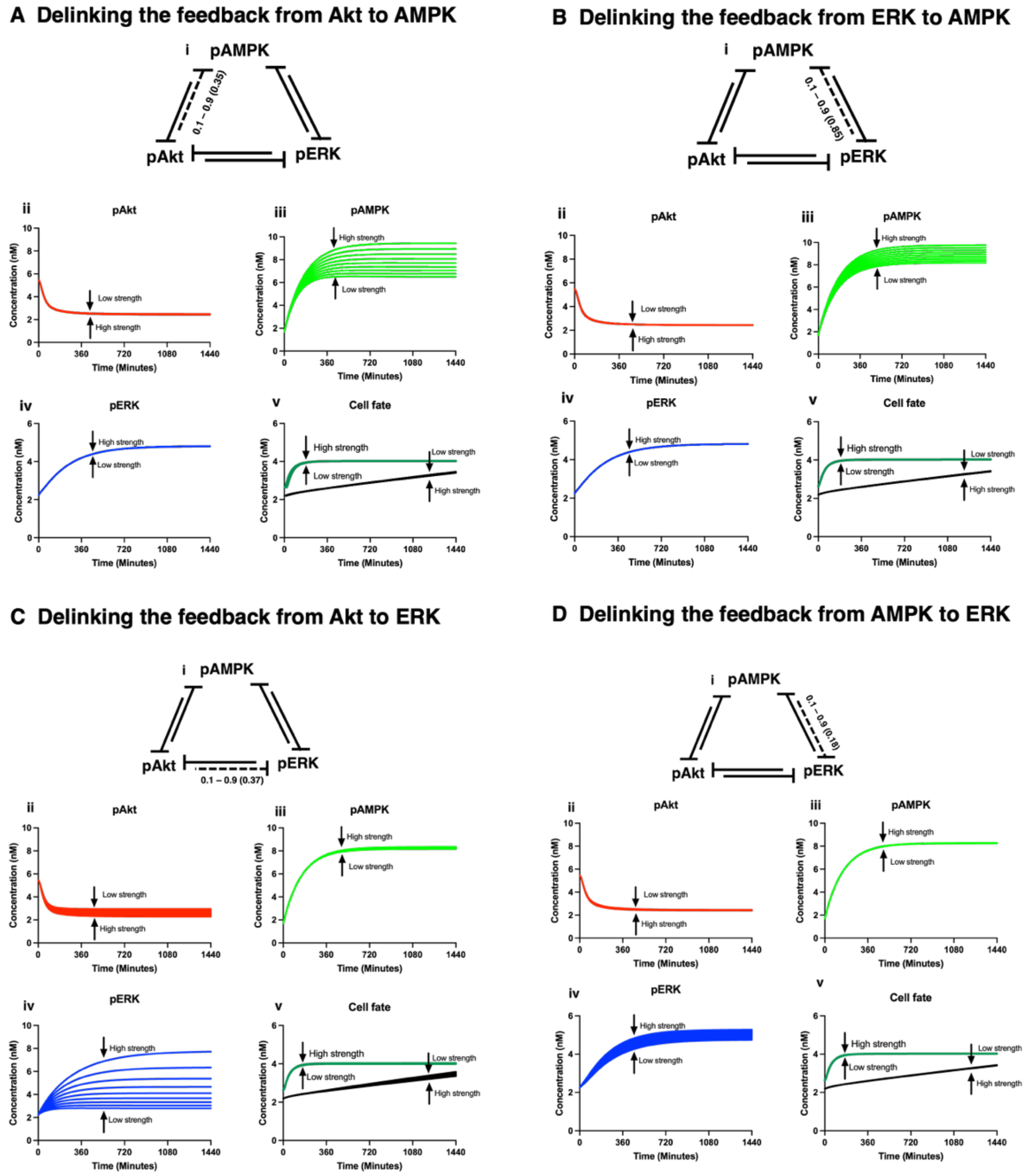


**Figure S11: The effect of feedback perturbation on pAkt, pAMPK, pERK and cell fate. (A**) Delinking the feedback from Akt to AMPK **(B)** Delinking the feedback from ERK to AMPK **(C)** Delinking the feedback from Akt to ERK **(D)** Delinking the feedback from AMPK to ERK.

**Table S1.** Summary of data used for model calibration and validation

| SR No. | Experimental condition | Attached to suspension | Timepoint of suspension | Method of detection | Range of experimental data | Reference |
|---|---|---|---|---|---|---|
| | **A. Calibration set** | | | | | |
| 1. | | **Levels of pAkt in sus vs. att** | | | | |
| | Cells subjected to matrix-deprivation and compared with attached cells | pAkt (sus)/pAkt(att) | 10 mins | WB | 0.4 - 0.5 | Supplementary Figure 3A |
| | | pAkt (sus)/pAkt(att) | 10 mins | WB | 0.4 - 0.5 | Saha et al., 2018 |
| | | pAkt (sus)/pAkt(att) | 8 h | WB | 0.4 - 0.5 | Supplementary Figure 3B |
| | | pAkt (sus)/pAkt(att) | 8 h | WB | 0.56 | Saha et al., 2018 |
| 2. | | **Levels of pAMPK in sus vs. att** | | | | |
| | | pAMPK (sus)/pAMPK(att) | 10 mins | WB | 4 - 5 | Supplementary Figure 3A |
| | | pAMPK (sus)/pAMPK(att) | 10 mins | WB | 1.5 - 2.5 | Saha et al., 2018 |
| | | pAMPK (sus)/pAMPK(att) | 8 h | WB | 6 - 7 | Supplementary Figure 3B |
| | | pAMPK (sus)/pAMPK(att) | 10 mins | WB | 4 - 5 | Sundararaman et al., 2016. |
| | | pAMPK (sus)/pAMPK(att) | 8 h | WB | 4 - 5 | Sundararaman et al., 2016. |
| | | pAMPK (sus)/pAMPK(att) | 24 h | WB | 5 - 6 | Sundararaman et al., 2016. |
| 3. | | **Levels of pERK in sus vs. att** | | | | |
| | | pERK(sus)/pERK(att) | 1 h | WB | 1.2 | Supplementary Figure 3C |
| | | pERK(sus)/pERK(att) | 2 h | WB | 0.71 | Supplementary Figure 3C |
| | | pERK(sus)/pERK(att) | 8 h | WB | 2 - 3 | Supplementary Figure 3C |
| | | pERK(sus)/pERK(att) | 24 h | WB | 4 - 5 | Supplementary Figure 3C |
| | | pERK(sus)/pERK(att) | 24 h | WB | 2 - 3 | Kumar et al., 2019, bioRxiv. |
| 4. | | **Levels of autophagy** | | | | |

| | | | | | | |
|---|---|---|---|---|---|---|
| | | Autophagosome(sus)/ autophagosome(att) | 24 h | Autophagosome detection by fluorescence microscopy | 1.5 – 2.0 | Figure 2H |
| 5. | | **Levels of apoptosis** | | | | |
| | | AnnexinV/PI(sus)/ AnnexinV/PI(att) | 24 h | AnnexinV/PI detection by flowcytometry | 1.4 – 1.7 | Figure 1A-B |
| | | **Attached condition** | | | | |
| | **AMPK activation (by genetic approaches) in adherent cells decreases pAkt levels** | pAkt (genetic approach)/pAkt (control) | - | WB | 0.4 – 0.5 | Supplementary Figure 4 |
| 8. | **ERK inhibition in attached cells increases pAkt levels** | pAkt (ERK inhibitor)/ pAkt (DMSO) | - | WB | 1.3 – 2.0 | Stulpinas et al., 2023 |
| 9. | **Akt inhibition in attached cells increases pERK levels** | pERK (Akt inhibitor)/ pERK (DMSO) | - | WB | 1.2 – 1.6 | Stulpinas et al., 2023 |
| | | **Suspension condition** | | | | |
| | **B. Validation** | | | | | |
| | | **Suspension condition** | | | | |
| 1. | **Akt hyperactivation enhances apoptosis in suspension cells** | Cleaved caspase3 (GFP Akt DD)/ Cleaved caspase3 (GFP) | 48 h | WB | 1.8 – 2.3 | Saha et al., 2018 |

| | | | | | | |
|---|---|---|---|---|---|---|
| **2.** | **Akt hyperactivation reduces number of anchorage-independent colonies in suspension** | Colonies (GFP Akt DD)/<br>colonies (GFP) | 15 days | Colony formation assay | 0.3 – 0.4 | Saha et al., 2018 |
| **3.** | **AMPK inhibition enhances apoptosis in suspension cells** | Caspase-3 activity (CC)/<br>Caspase-3 activity (DMSO) | 48 h | WB | 1.6 – 2 | Saha et al., 2018 |
| **4.** | **AMPK inhibition reduces number of anchorage-independent colonies in suspension** | Colonies (CC)/<br>colonies (CC) | 15 days | WB | 0.1 – 0.2 | Saha et al., 2018 |
| **5.** | **Down-regulation of PHLPP2 leads to increased pAkt in suspension cells** | pAkt (PHLPP2 KD)/<br>pAkt(control) | 8 h | WB | 3 – 7 | Supplementary Figure 6A |
| **6.** | **Down-regulation of PHLPP2 leads to decreased pAMPK in suspension cells** | pAMPK (PHLPP2 KD)/<br>pAMPK(control) | 8 h | WB | 0.8 – 0.9 | Supplementary Figure 6A |
| **7.** | **Akt inhibition in PHLPP2 KD cells lead to** | pAkt (PHLPP2 KD)/<br>pAkt (control) | 48 h | WB | 0.2 – 0.4 | Supplementary Figure 6B |

| | | | | | | |
|---|---|---|---|---|---|---|
| | **decreased pAkt in suspension** | | | | | |
| 8. | **PHLPP2 KD enhances apoptosis in suspension cells** | Caspase-3 activity (PHLPP2 KD)/ Caspase-3 activity (control) | 48 h | Caspase-3 activity assay | 3.5 – 4.5 | Saha et al., 2018 |
| 9. | **PHLPP2 KD reduces number of anchorage-independent colonies in suspension** | Colonies (PHLPP2 KD)/ colonies (PHLPP2 KD) | 48 h | Soft agar colony formation assay | 0.2 – 0.4 | Saha et al., 2018 |
| 10. | **Akt inhibition in PHLPP2 KD cells lead to increased pAMPK in suspension** | pAMPK (PHLPP2 KD)/ pAMPK (control) | 48 h | WB | 1 - 3 | Supplementary Figure 6B |

**Table S2.** List of variables (molecular players) in the anoikis resistance model

| Sr. No. | Protein species | Description |
|---|---|---|
| 1 | **Matrix (input)** | Status of matrix attachment or detachment |
| 2 | **Ccal** | Cytosolic calcium |
| 3 | tAMPK | AMP-activated protein kinase |
| 4 | **pAMPK** | Phosphorylated AMP-activated protein kinase |
| 5 | tAkt | Protein kinase B |
| 6 | **pAkt** | Phosphorylated protein kinase B |
| 7 | tERK | Extracellular Signal-Regulated kinase |
| 8 | **pERK** | Phosphorylated Extracellular Signal-Regulated kinase |
| 9 | proCaspase | Pro-caspase |
| 10 | **aCaspase** | Active-caspase |
| 11 | **Phagophore** | Phagophore |

| | | |
|---|---|---|
| 12 | **Autophagosome** | Autophagosome |

#### Model equations

The equations below remain the same across matrix-attached and matrix-deprived state. Based on the levels of the proteins in their respective states, they exert either excitatory or inhibitory effects.

##### 1. Equation for Akt

$$\frac{dpAkt}{dt} = \text{activation of Akt} - \text{ deactivation of pAkt influenced by pAMPK and pERK}$$

$$\frac{dpAkt}{dt} = k_a Akt * (tAkt - pAkt) - k_d Akt * pAkt * V_{1a} * \left(\frac{pAMPK^{n1a}}{pAMPK^{n1a} + k_{1a}^{n1a}}\right) * V_{1b} * \left(\frac{pERK^{n1b}}{pERK^{n1b} + k_{1b}^{n1a}}\right)$$

Description: In the matrix-attached state, Akt is activated via growth factor and integrin signaling. The details of growth factor and integrin signaling are avoided for model simplicity. In the matrix-deprived state, both AMPK and ERK are active and they exert an inhibitory effect on Akt. Since both AMPK and ERK exert effects on Akt in the indirect manner, there inhibitory effects have been included as Hill functions.

##### 2. Equation for AMPK

$$\frac{dpAMPK}{dt} = \text{activation of AMPK influenced by CAMKK}\beta - \text{ deactivation of pAMPK influenced by pAkt and pERK}$$

$$\frac{dpAMPK}{dt} = k_a AMPK * CAMKK\beta * (tAMPK - pAMPK) - k_d AMPK * pAMPK * V_{2a} * \left(\frac{pAkt^{n2a}}{pAkt^{n2a} + k_{2a}^{n2a}}\right) * V_{2b} * \left(\frac{pERK^{n2b}}{pERK^{n2b} + k_{2b}^{n2b}}\right)$$

Description: In the matrix-deprived state, there is a spike in cytosolic calcium which activates AMPK via CAMKKβ. Matrix-deprived state also activates ERK. Active ERK (pERK) exerts an inhibitory effect on pAMPK. Though levels of pAkt are maintained low in matrix-detached state, pAkt also exerts an inhibitory effect on pAMPK.

##### 3. Equation for ERK

$$\frac{dpERK}{dt} = \text{activation of ERK influenced by matrix deprivation } + \text{ activation of ERK hindered by pAMPK} - \mathit{deactivation\ of\ pERK\ influenced\ by\ pAkt}$$

$$\frac{dpERK}{dt} = k_a ERK * (2 - Matrix) * (tERK - pERK) + k_a ERK * V_{3a} * \left(\frac{k_{3a}^{n3a}}{pAMPK^{n3a} + k_{3a}^{n3a}}\right) * (tERK - pERK) - k_d ERK * pERK * V_{3b} * \left(\frac{pAkt^{3b}}{pAkt^{3b} + k_{3b}^{n3b}}\right)$$

Description: Activation of ERK is facilitated by matrix deprivation but it is hindered by pAMPK. pAMPK phosphorylates PEA15 in matrix-deprived cells and hinders MEK-mediated ERK phosphorylation. Hence, AMPK prevents the phosphorylation of ERK indirectly. Though levels of pAkt are maintained low in matrix-detached state, pAkt exerts an inhibitory effect on pERK.

4. **Equation for aCaspase (Apoptosis)**

$$\frac{\mathrm{daCaspase}}{\mathrm{dt}} = \text{Basal activation of caspases} + \text{activation of caspases influenced by pERK} + \text{activation of caspases influenced by pAKt specifically in matrix} - \text{deprived state} + \text{activation of caspases hindered by pAMPK}$$

$$\frac{\mathrm{daCaspase}}{\mathrm{dt}} = \mathrm{k_aCas_{Basal}} * (\mathrm{proCaspase} - \mathrm{aCaspase}) + \mathrm{k_aCas} * \mathrm{V_{4a}} * \left(\frac{\mathrm{pERK^{n4a}}}{\mathrm{pERK^{n4a}} + \mathrm{k_{4a}^{n4a}}}\right) * (\mathrm{proCaspase} - \mathrm{aCaspase}) + \mathrm{k_aCas} * (12 - 12 * \mathrm{Matrix}) * \mathrm{V_{4b}} * \left(\frac{\mathrm{pAkt^{n4b}}}{\mathrm{pAkt^{n4b}} + \mathrm{k_{4b}^{n4b}}}\right) * (\mathrm{proCaspase} - \mathrm{aCaspase}) + \mathrm{k_aCas} * \mathrm{V_{4c}} * \left(\frac{\mathrm{k_{4c}^{n4c}}}{\mathrm{pAMPK^{n4c}} + \mathrm{k_{4c}^{n4c}}}\right) * (\mathrm{proCaspase} - \mathrm{aCaspase})$$

Description: The dynamics of apoptosis are captured by the activation of caspases. Once caspases are activated, a cell is programmed to undergo cell death via the process of apoptosis. pERK and pAkt aids in the activation of caspases, whereas pAMPK hinders the activation of caspases. Caspases do not undergo any deactivation.

5. **Equation for phagophore**

$$\frac{\mathrm{dPhagophore}}{\mathrm{dt}} = \text{Basal rate of formation of phagophore} + \text{formation of phagophore facilitated by the stress of matrix deprivation} - \textit{dissociation of phagophore}$$

$$\frac{\mathrm{dPhagophore}}{\mathrm{dt}} = \mathrm{k_{f_{Pg\_Basal}}} + \mathrm{k_{f_{Pg}}} * (2 - \mathrm{Matrix}) - \mathrm{k_{d_{Pg}}} * \mathrm{Phagophore}$$

Description: The process of autophagy is facilitated by the formation of autophagosome from phagophores. Since phagophores are dynamic double-membrane structures, it is included in the model as a differential equations instead of a constant. Phagophore formation is facilitated due to the stress of matrix deprivation, in addition to the basal rate of formation and dissociation.

6. **Equation for autophagosome (Autophagy)**

$$\frac{dAutophagosome}{dt} = \text{Basal rate of formation of autophagosome} + \text{formation of autophagosome facililated by pAMPK} + \text{formation of autophagosome hindered by pAkt} + \text{formation of autophagome hindered by pERK} - \text{fusion of autophagosome with lysosome and degradation}$$

$$\frac{dAutophagosome}{dt} = k_f Ag * (Phagophore - Autophagosome) + k_f Ag * V_{5a} * \left(\frac{pAMPK^{n5a}}{pAMPK^{n5a} + k_{5a}^{n5a}}\right) * (Phagophore - Autophagosome) + k_f Ag * V_{5b} * \left(\frac{k_{5a}^{n5b}}{pAkt^{n5b} + k_{5a}^{n5b}}\right) * (Phagophore - Autophagosome) + k_f Ag * V_{5c} * \left(\frac{k_{5c}^{n5c}}{pERK_{5c}^{n5c} + k_{5c}^{n5c}}\right) * (Phagophore - Autophagosome) - k_d Ag * Autophagosome$$

Description: Basal autophagy facilitates the recycling of nutrients. In addition to basal autophagy, it is facilitated by pAMPK and hindered by pAkt and pERK. Once the autophagosome fuses with lysosome, its contents are degraded with the help of the lysosomal enzymes.

#### Table S3. Initial conditions of all variables (molecular players)

| Sr. No. | Protein species | Initial conditions | Steady state value (Matrix-attached state) | Steady state value (Matrix-deprived state) |
|---|---|---|---|---|
| 1 | Matrix (input) | 1 (Matrix-attached state)<br>0 (Matrix-deprived state) | 1 | 0 |
| 2 | Ccal | 1 (Matrix-attached state)<br>2 (Matrix-deprived state) | 1 | 2 |
| 3 | tAMPK | 10 | 10 | 10 |
| 4 | pAMPK | 0 | 1.70 | 8.26 |
| 5 | tAkt | 10 | 10 | 10 |
| 6 | pAkt | 0 | 5.44 | 2.44 |
| 7 | tERK | 10 | 10 | 10 |

| | | | | |
|---|---|---|---|---|
| 8 | pERK | 0 | 2.25 | 4.81 |
| 9 | proCaspase | 10 | 10 | 10 |
| 10 | aCaspase | 0 | 3.07 | 3.42 |
| 11 | Phagophore | 0 | 2.98 | 4.12 |
| 12 | Autophagosome | 0 | 2.63 | 4.02 |

#### Table S4. List of parameter values

| Sr. No. | Parameter values | Description | Values | Units | References |
|---|---|---|---|---|---|
| 1 | **Dynamics of Akt** | | | | |
| | $k_a Akt$ | Rate of activation of Akt | 0.0189 | $nM^{-1}min^{-1}$ | Liu et al., 2017 |
| | $k_d Akt$ | Rate of deactivation of pAkt | 0.0176 | $min^{-1}$ | Liu et al., 2017 |
| | $V_{1a}$ | Fold change of Akt inhibition by pAMPK | 2 | Dimensionless | Calibrated |
| | $n_{1a}$ | Hill coefficient | 2.5 | Dimensionless | Calibrated |
| | $k_{1a}$ | Threshold of Akt inhibition by pAMPK | 2 | nM | Calibrated |
| | $V_{1b}$ | Fold change of Akt inhibition by pERK | 2 | Dimensionless | Calibrated |
| | $n_{1b}$ | Hill coefficient | 2 | Dimensionless | Calibrated |
| | $k_{1b}$ | Threshold of Akt inhibition by pERK | 2 | nM | Calibrated |
| 2. | **Dynamics of AMPK** | | | | |
| | $k_a AMPK$ | Rate of activation of AMPK | 0.000164 | $nM^{-1}min^{-1}$ | Liu et al., 2017 |
| | $k_d AMPK$ | Rate of deactivation of pAMPK | 0.0159 | $min^{-1}$ | Liu et al., 2017 |

| | | | | | |
|---|---|---|---|---|---|
| | $V_{Ccal}$ | Fold change of CAMKKβ activation by Ccal | 37 | Dimensionless | Maiti et al., 2024 |
| | $n_{Ccal}$ | Hill coefficient | 8 | Dimensionless | Calibrated |
| | $k_{Ccal}$ | Threshold of CAMKKβ activation by Ccal | 1.5 | nM | Calibrated |
| | $V_{2a}$ | Fold change of AMPK inhibition by pAkt | 0.5 | Dimensionless | Calibrated |
| | $n_{2a}$ | Hill coefficient | 3 | Dimensionless | Calibrated |
| | $k_{2a}$ | Threshold of AMPK inhibition by pAkt | 3 | nM | Calibrated |
| | $V_{2b}$ | Fold change of AMPK inhibition by pERK | 0.5 | Dimensionless | Calibrated |
| | $n_{2b}$ | Hill coefficient | 2 | Dimensionless | Calibrated |
| | $k_{2b}$ | Threshold of AMPK inhibition by pERK | 2 | nM | Calibrated |
| **3.** | **Dynamics of ERK** | | | | |
| | $k_a ERK$ | Rate of activation of ERK | 0.0012 | $nM^{-1}min^{-1}$ | Aoki et al., 2011 |
| | $k_d ERK$ | Rate of deactivation of pERK | 0.0024 | $min^{-1}$ | Todd et al., 1999 |
| | $V_{3a}$ | Fold change of ERK inhibition by pAMPK | 0.5 | Dimensionless | Calibrated |
| | $n_{3a}$ | Hill coefficient | 2 | Dimensionless | Calibrated |
| | $k_{3a}$ | Threshold of ERK inhibition by pAMPK | 4 | nM | Calibrated |
| | $V_{3b}$ | Fold change of ERK inhibition by pAkt | 3 | Dimensionless | Calibrated |
| | $n_{3b}$ | Hill coefficient | 2.5 | Dimensionless | Calibrated |

| | | | | | |
|---|---|---|---|---|---|
| | $k_{3b}$ | Threshold of ERK inhibition by pAkt | 3 | nM | Calibrated |
| **4.** | **Dynamics of Caspase** | | | | |
| | $k_a Cas_{Basal}$ | Basal rate of activation of caspases | 1.67 x $10^{-6}$ | $nM^{-1}min^{-1}$ | Calibrated |
| | $k_a Cas$ | Rate of activation of caspases | 9 x $10^{-6}$ | $nM^{-1}min^{-1}$ | Calibrated |
| | $V_{4a}$ | Fold change of caspase activation by pERK | 2 | Dimensionless | Calibrated |
| | $n_{4a}$ | Hill coefficient | 3 | Dimensionless | Calibrated |
| | $k_{4a}$ | Threshold of caspase activation by pERK | 3 | nM | Calibrated |
| | $V_{4b}$ | Fold change of caspase activation by pAkt | 3 | Dimensionless | Calibrated |
| | $n_{4b}$ | Hill coefficient | 2 | Dimensionless | Calibrated |
| | $k_{4b}$ | Threshold of caspase activation by pAkt | 4 | nM | Calibrated |
| | $V_{4c}$ | Fold change of hindrance to caspase activation by pAMPK | 10 | Dimensionless | Calibrated |
| | $n_{4c}$ | Hill coefficient | 3 | Dimensionless | Calibrated |
| | $k_{4c}$ | Threshold of hindrance to caspase activation by pAMPK | 3 | nM | Calibrated |
| **5.** | **Dynamics of Autophagosome** | | | | |
| | $k_{f_{Pg_{Basal}}}$ | Basal rate of formation of phagophore | 0.0167*2 | $nM^{-1}min^{-1}$ | Calibrated |

| | | | | | |
|---|---|---|---|---|---|
| | $k_{f_{Pg}}$ | Rate of formation of phagophore | 0.017*1.2 | $nM^{-1}min^{-1}$ | Calibrated |
| | $k_{d_{Pg}}$ | Rate of dissociation of phagophore | 0.018 | $min^{-1}$ | Liu et al., 2017 |
| | $k_{f_{Ag_{Basal}}}$ | Basal rate of formation of autophagosome | 0.0167 | $nM^{-1}min^{-1}$ | Liu et al., 2017 |
| | $k_{f_{Ag}}$ | Rate of formation of autophagosome | 0.0167 | $nM^{-1}min^{-1}$ | Liu et al., 2017 |
| | $k_{d_{Ag}}$ | Rate of degradation of autophagosome | 0.0167 | $min^{-1}$ | Liu et al., 2017 |
| | $V_{5a}$ | Fold change of autophagosome formation activated by pAMPK | 5 | Dimensionless | Calibrated |
| | $n_{5a}$ | Hill coefficient | 4 | Dimensionless | Calibrated |
| | $k_{5a}$ | Threshold of autophagosome formation activated by pAMPK | 2 | nM | Calibrated |
| | $V_{5b}$ | Fold change of autophagosome formation inhibited by pAkt | 100 | Dimensionless | Calibrated |
| | $n_{5b}$ | Hill coefficient | 3 | Dimensionless | Calibrated |
| | $k_{5b}$ | Threshold of autophagosome formation inhibited by pAkt | 2 | nM | Calibrated |
| | $V_{5c}$ | Fold change of autophagosome | 1 | Dimensionless | Calibrated |

| | | | | | |
|---|---|---|---|---|---|
| | | formation inhibited by pERK | | | |
| | $n_{5c}$ | Hill coefficient | 2 | Dimensionless | Calibrated |
| | $k_{5c}$ | Threshold of autophagosome formation inhibited by pERK | 0.5 | nM | Calibrated |

**MATLAB Code**

Function file

```
function H1 = AR(t, H);
pAkt           = H(1);
pAMPK          = H(2);
pERK           = H(3);
aCaspase       = H(4);
Phagophore     = H(5);
Autophagosome = H(6);
Ccal           = H(7);
Matrix         = H(8);
%% Matrix
Matrixprime = 0;
%% Ccal
Ccalprime = 0;
%% 1. Akt
ka_akt = 0.0189;  %  0.0189 nM-1min-1
kd_akt = 0.0176;  %  min-1;   (Akt inhibition: 0.3 -
0.7, *2.15 gives 1.62), Akt inhibition on PHPLL2 KD:
*100, *500
%GF      = 1;
tAkt   = 10;
V1A    = 2; % 10, 7, 5
n1A    = 2.5;
k1A    = 2; %3
Hf1A   = V1A * (pAMPK^n1A/(pAMPK^n1A + k1A^n1A)); %
PHLPP2 KD: 0.01
V1B    = 2;%3
n1B    = 2; % 2,3,4
k1B    = 2; % 7,5
Hf1B   = V1B * (pERK^n1B/(pERK^n1B + k1B^n1B));
```

```
pAktprime = (ka_akt * (tAkt - pAkt)) - (kd_akt * pAkt *
Hf1A * Hf1B);
%% 2. AMPK
ka_ampk = 0.000164; % nM-1min-1 (0.000164 * 4.5) %
PHLPP2 KD - ampk activation *5
kd_ampk = 0.0159; % min-1, AMPK inhibition: *5 and *10
tAMPK   = 10;% nM
VCcal = 37;
nCcal = 8;
kCcal = 1.5;
CAMKKB  = 1 + VCcal *(Ccal^nCcal / (Ccal^nCcal +
kCcal^nCcal));
V2A     = 0.5; %0.5
n2A     = 3;   % 3 (Final)
k2A     = 3;   %0.5
Hf2A    = V2A * (pAkt^n2A/(pAkt^n2A + k2A^n2A));
V2B     = 0.5;
n2B     = 2;
k2B     = 2;%2, 0.5
Hf2B    = V2B * (pERK^n2B/(pERK^n2B + k2B^n2B));
% Hf2B    = V2B * (k2B^n2B/(pERK^n2B + k2B^n2B));
% pAMPKprime = (ka_ampk * CAMKKB * (tAMPK - pAMPK)) -
(kd_ampk * pAMPK * Hf2A * Hf2B);
pAMPKprime = (ka_ampk * CAMKKB  * (tAMPK - pAMPK)) -
(kd_ampk * pAMPK * Hf2A * Hf2B);
%% 3. pERK
ka_ERK  = 0.0012;         % nM-1min-1
kd_ERK  = 0.0024; % min-1 (0.0024 * 3)
tERK    = 10;
V3A     = 0.5; %1
n3A     = 2;
k3A     = 4;%1, 0.25
Hf3A    = V3A * (k3A^n3A/(pAMPK^n3A + k3A^n3A));
V3B     = 3; % 1
n3B     = 2.5;%1
k3B     = 3; %8
Hf3B    = V3B *(pAkt^n3B/(k3B^n3B + pAkt^n3B)); % Akt's
inhibition on ERK delink:  V3B * 0.01;
pERKprime = ka_ERK * Hf3A * (tERK - pERK) + ka_ERK * (2
- Matrix) * (tERK - pERK) - kd_ERK * pERK * Hf3B;
%% 4. aCaspase
ka_cas_basal = 0.00000167;
ka_cas    = 0.000009;  % nM-1min-1
kd_cas    = 0.000000167;  % 0.0165 nM-1min-1
```

```
proCaspase = 10;
V4A      = 2;
n4A      = 3;
k4A      = 3; %4
Hf4A     = V4A * (pERK^n4A/(pERK^n4A + k4A^n4A));
V4B      = 3;
n4B      = 2;
k4B      = 4; %1
Hf4B     = V4B * (pAkt^n4B/(pAkt^n4B + k4B^n4B));
V4C      = 10; % 0.5
n4C      = 3; % 2.5
k4C      = 3; %8
Hf4C     = V4C * (k4C^n4C/(pAMPK^n4C + k4C^n4C));
aCasprime = ka_cas_basal * (proCaspase - aCaspase) +
ka_cas * Hf4A * (proCaspase - aCaspase) + ka_cas * (12
- (12 * Matrix)) * Hf4B * (proCaspase - aCaspase) +
ka_cas * Hf4C * (proCaspase - aCaspase);%-  kd_cas *
aCaspase;
%% 5. Autophagosome
kf_Ag_Basal  = 0.0167;
kf_Ag  = 0.0167;  % nM-1min-1
kd_Ag  = 0.0167;  % nM-1min-1
% Phagophore = 10;
V5A     = 5; %5
n5A     = 4;
k5A     = 2;%2
Hf5A    = V5A * (pAMPK^n5A/(pAMPK^n5A + k5A^n5A));
V5B     = 100; %0.1
n5B     = 3;%3
k5B     = 2;%2
Hf5B    = V5B * (k5B^n5B/(pAkt^n5B + k5B^n5B));
V5C     = 1; % 0.5
n5C     = 2;
k5C     = 0.5;
Hf5C    = V5C * (k5C^n5C/(pERK^n5C + k5C^n5C));
kf_Pg_basal = 0.0167*2;
kf_Pg = 0.017*1.2;
kd_Pg = 0.018;%*0.3
Phagophoreprime = kf_Pg_basal + kf_Pg * (2 - Matrix) -
kd_Pg * Phagophore;
Autophagosomeprime = kf_Ag * (Phagophore -
Autophagosome) + kf_Ag * Hf5A * (Phagophore -
Autophagosome) + kf_Ag * Hf5B * (Phagophore -
```

```
Autophagosome) + kf_Ag * Hf5C * (Phagophore -
Autophagosome) - kd_Ag * Autophagosome;
%% Outputs
H1 = [pAktprime; pAMPKprime; pERKprime;
aCasprime;Phagophoreprime; Autophagosomeprime;
Ccalprime; Matrixprime;];
```

Run file:

```
clc
clear all
close all
% k = [ 0 0];
 % 1 (attached), 0 (suspension)
 % IC = 1:-0.01:0; % att to sus
%  IC = 0:0.01:1;
%  NC = (zeros(length(IC),8));
% for i = 1: length (IC)
% Matrix = IC(i);

Matrix = 1; % 1(natrix-attached state), 0(matrix-
deprived state)
x = 2;
Ccal = x - Matrix;
 % k = [0 0 0 0 0 0 Ccal Matrix];
 % Knockdown studies 0.001
 % pAMPK IC att = 1.70433330976211
 % pAkt IC att  = 5.44891414031187
 % pERK IC att  = 2.25097027832944, KD=0.001
 % att ss
%  IC = 0.1:0.1:10;
% NC = (zeros(length(IC),8));
% for i = 1: length (IC)
% att ss
% k = [0 0 0 0 0 0 Ccal Matrix];% sus ss 60 mins

  k = [5.44891414031187  1.70433330976211
2.25097027832944 2.19541246480736  2.98888887227600
    2.63662952975674  Ccal Matrix];% sus ss 60 mins
% sus ss 60 mins
% k = [3.68339472581702    3.82929598747606
    2.72327480340665  2.30723962311656
    3.73718103923567  3.52741804355827 Ccal Matrix];
% sus ss 1440 mins
```

```
  % k = [2.44637624561553  8.26446960540757
     4.81461775952763  3.42517438097746
     4.12222157994473  4.02483329123111  Ccal Matrix];
 % k = [5.63145850424296   1.64020252572228
     2.22444569424266  1.19952231477569
     2.06111073684133  1.49802523230363 Ccal Matrix];
 % k = [5.63173213901539   1.64019236227446
     2.22449455699597  1.19952306602864
     4.23333324423632  3.07672371891565  Ccal Matrix];
 % k=[5.631732139015391.64019236227446
     2.22449455699597  1.19952306602864
     4.23333324423632  3.07672371891565 Ccal Matrix];
% k = [5.63158148386812    1.64016620474462
     2.22445550564726  2.98528134631546
     4.23333310051570  3.07679301473951 Ccal Matrix];
  % k = [5.63080036011786  1.64015787931123
     2.22426600488555  1.18059653639280
     3.14669268460282  2.33890879473237  Ccal Matrix];
% k = [5.63089438266742    1.64041752771945
     2.22454526925774  1.73586061081245
     1.25795715504609  Ccal Matrix];
% k = [5.63149253796546    1.64020614872363
     2.22445356915330  0.170343244996904
     1.25753397300259 Ccal Matrix];
% k = [5.63122269625512    1.64029601595078
     2.22453593725755  1.08053732993395  Ccal Matrix];
% k = [2.44642411247972    8.26422719507496
     4.81417877316854  Ccal Matrix]; % 24h SS value at
sus
% k = [5.63158620932352    1.64015244012352
     2.22445072894411 0    Ccal Matrix]; % 24h SS value
at att


tspan = 0:1440;
H0 = k;
[t, H] = ode15s(@ARfunc, tspan, H0);
 % NC(i,:)= H(end,1:8);
 % end

%% Automating loops
```

```
% k = [5.63158620932352    1.64015244012352
    2.22445072894411  Ccal Matrix];

%storing values
tic;
pAkt          = H(:,1);
pAMPK         = H(:,2);
pERK          = H(:,3);
aCaspase      = H(:,4);
Phagophore    = H(:,5);
Autophagosome = H(:,6);
Ccal          = H(:,7);
Matrix        = H(:,8);
toc;

%% Calculating the net metabolic state as a function of
AMPK (catabolic) and Akt (anabolic)

 % AMPKmax = 9.05;
 % AMPKbs  = 1.88;
 % h_AMPK=(0.6.*(AMPKmax - AMPKbs));
 % Aktmax  = 5.15;
 % Aktbs  = 1.80;
 % h_Akt=(0.6.*(Aktmax - Aktbs));
 % Phst_C = pAMPK ./(h_AMPK + pAMPK);
 % Phst_A = pAkt ./(h_Akt + pAkt);
 %% Defining variables

 SS_Autophagosome = H(end,6);
 SS_aCaspase = H(end,4);
 %% Autophagy module
 Autophagy = SS_Autophagosome;
 %% Apoptosis module
 Apoptosis = SS_aCaspase;
 %% Cell survival
 Cell_survival = Autophagy - Apoptosis;
 %% Print Autophagy
V5A     = 5; %5
n5A     = 4;
k5A     = 2;%2
Hf5A    = V5A * (pAMPK.^n5A./(pAMPK.^n5A + k5A.^n5A));
V5B     = 0.1;%2 %0.1
n5B     = 3;%3
k5B     = 2;%2
```

```
Hf5B    = V5B * (k5B.^n5B./(pAkt.^n5B + k5B.^n5B));
V5C     = 0.5; % 0.5
n5C     = 2;
k5C     = 0.5;
Hf5C    = V5C * (k5C.^n5C./(pERK.^n5C + k5C.^n5C));
Ran = 1/(2 + Hf5A + Hf5B + Hf5C);
 %% Outputs

 plot(t, H(:,1), 'r', 'linewidth', 4)
 hold on;
 plot(t, H(:,2), 'g', 'linewidth', 4)
 hold on;
 plot(t, H(:,3), 'b', 'linewidth', 4)
 hold on;
 plot(t, H(:,4), 'c', 'linewidth', 4)
 hold on;
 plot(t, H(:,5), 'm', 'linewidth', 4)
 hold on;
 plot(t, H(:,6), 'k', 'linewidth', 4)
 hold on;
 % plot(t, H(:,7),'c','linewidth', 2)
 % hold on;


 % plot(t, Phst_A, 'b', 'linewidth', 2);
 % plot(t, Phst_C, 'm', 'linewidth', 2);


  legend("pAkt" ,"pAMPK", "pERK", "aCaspase3",
"Phagophore","Autophagosome");

 % legend("Anabolic state" ,"Catabolic state");

% legend("pAkt" ,"pAMPK");
ylabel("Concentration (nM)")
xlabel("Time (min)")
axis([0 1440 0 10])

%output = cat(pAkt,pAMPK_IC, pERK);
%end
```
